# Profiling of accessible chromatin regions across multiple plant species and cell types reveals common gene regulatory principles and new control modules

**DOI:** 10.1101/167932

**Authors:** Kelsey A. Maher, Marko Bajic, Kaisa Kajala, Mauricio Reynoso, Germain Pauluzzi, Donnelly A. West, Kristina Zumstein, Margaret Woodhouse, Kerry Bubb, Michael W. Dorrity, Christine Queitsch, Julia Bailey-Serres, Neelima Sinha, Siobhan M. Brady, Roger B. Deal

**Affiliations:** Department of Biology, Emory University, Atlanta, GA 30322; Graduate Program in Biochemistry, Cell, and Developmental Biology, Emory University, Atlanta, GA 30322; Graduate Program in Genetics and Molecular Biology, Emory University, Atlanta, GA 30322; Department of Plant Biology and Genome Center, University of California, Davis, CA 95616; Center for Plant Cell Biology, Botany and Plant Sciences Department, University of California, Riverside, Riverside, CA 92521; Department of Plant Biology, University of California, Davis, CA 95616; University of Washington, School of Medicine, Department of Genome Sciences, Seattle, WA 98195; Present Address: Plant Ecophysiology, Institute of Environmental Biology, Utrecht University 3584 CH, Utrecht, the Netherlands

**Author notes:** These authors contributed equally to this work (Kelsey A. Maher, Marko Bajic). Correspondence: Roger B. Deal.

## Abstract

The transcriptional regulatory structure of plant genomes remains poorly defined relative to animals. It is unclear how many *cis*-regulatory elements exist, where these elements lie relative to promoters, and how these features are conserved across plant species. We employed the Assay for Transposase-Accessible Chromatin (ATAC-seq) in four plant species (*Arabidopsis thaliana*, *Medicago truncatula*, *Solanum lycopersicum*, and *Oryza sativa*) to delineate open chromatin regions and transcription factor (TF) binding sites across each genome. Despite 10-fold variation in intergenic space among species, the majority of open chromatin regions lie within 3 kb upstream of a transcription start site in all species. We find a common set of four TFs that appear to regulate conserved gene sets in the root tips of all four species, suggesting that TF-gene networks are generally conserved. Comparative ATAC-seq profiling of *Arabidopsis* root hair and non-hair cell types revealed extensive similarity as well as many cell type-specific differences. Analyzing TF binding sites in differentially accessible regions identified a MYB-driven regulatory module unique to the hair cell, which appears to control both cell fate regulators and abiotic stress responses. Our analyses revealed common regulatory principles among species and shed light on the mechanisms producing cell type-specific transcriptomes during development.

## INTRODUCTION

The transcription of protein coding genes is controlled by regulatory DNA elements, including both the core promoter and more distal enhancer elements (Lee and Young, 2000). The core promoter is a short DNA region surrounding the transcription start site (TSS), at which RNA polymerase II and general transcription factors are recruited. Enhancer elements act as platforms for recruiting both positive- and negative-acting transcription factors (TFs), and serve to integrate multiple signaling inputs in order to dictate the spatial and temporal control of transcription from the core promoter. As such, enhancer functions are critical for directing transcriptional output during cell differentiation and development, as well as coordinating transcriptional responses to environmental change (Ong and Corces, 2011). Despite their importance, only a small number of *bona fide* enhancers have been characterized in plants, and we lack a global view of their general distribution and action in plant genomes (Weber et al., 2016).

In large part, our limited knowledge of plant *cis*-regulatory elements arises from the unique difficulties in identifying these elements. While some enhancers exist near their target core promoter, others can be thousands of base pairs upstream or downstream, or even within the transcribed region of a gene body (Ong and Corces, 2011; Spitz and Furlong, 2012). Furthermore, enhancers generally do not display universal sequence conservation, aside from sharing of individual TF binding sites, which makes them very challenging to locate. By contrast, core promoters can be readily identified through mapping the 5’ ends of transcripts (Morton et al., 2014; Mejia-Guerra et al., 2015). It was recently discovered that many enhancer elements in animal genomes could be identified with relatively high confidence based on a unique combination of flanking histone posttranslational modifications (PTMs), such as an enrichment for H3K27ac and H3K4me1. This characteristic histone PTM signature has led to the annotation of such elements in several animal models and specialized cell types (Heintzman et al., 2009; Bonn et al., 2012). However, the only currently known association between plant *cis*-regulatory elements and histone PTMs appears to be a modest correlation with H3K27me3 (Zhang et al., 2012b; Zhu et al., 2015). Though encouraging, this mark is not unique to these elements, and cannot be used to identify enhancers on its own.

A long-known and general feature of sequence-specific DNA-binding proteins is their ability to displace nucleosomes upon DNA binding, leading to an increase in nuclease accessibility around the binding region (Gross and Garrard, 1988; Henikoff, 2008). In particular, DNaseI treatment of nuclei coupled with high-throughput sequencing (DNase-seq) has been used to probe chromatin accessibility. This technology has served as an important tool in identifying regulatory elements throughout animal genomes (Thurman et al., 2012) and more recently in certain plant genomes (Zhang et al., 2012b; Zhang et al., 2012a; Pajoro et al., 2014; Sullivan et al., 2014). In addition, a differential micrococcal nuclease sensitivity assay has also been used to probe functional regions of the maize genome, demonstrating the versatility of this approach (Vera et al., 2014; Rodgers-Melnick et al., 2016).

DNase-seq has been used successfully to identify open chromatin regions in different tissues of both rice and *Arabidopsis* (Zhang et al., 2012a; Pajoro et al., 2014; Zhu et al., 2015). Over a dozen of the intergenic DNase-hypersensitive sites in *Arabidopsis* were tested and shown to act as enhancer elements by activating a minimal promoter-reporter cassette, demonstrating that chromatin accessibility is an important factor in enhancer identification (Zhu et al., 2015). Collectively, these DNase-seq studies show that the majority of open chromatin sites exist outside of genes in rice and *Arabidopsis*, that differences in open chromatin sites can be identified between tissues, and that a large proportion of intergenic open chromatin sites are in fact regulatory, at least in *Arabidopsis*. Another recent significant advance came from using DNase-seq to examine the changes in *Arabidopsis* chromatin accessibility and TF occupancy that occur during development and in response to abiotic stress (Sullivan et al., 2014). This work showed that TF-to-TF regulatory network connectivity appears to be similar between *Arabidopsis*, human, and *C*. *elegans*, and that such networks were extensively ‘rewired’ in response to stress. This study also showed that many genetic variants linked to complex traits were preferentially located in accessible chromatin regions, portending the potential for harnessing natural variation in regulatory DNA for plant breeding.

We are still left with many open questions regarding the general conservation of transcriptional regulatory landscapes across plant genomes. For example, it remains unclear how many *cis*-regulatory elements generally exist in plant genomes, where they reside in relation to their target genes, and to what extent these features are conserved across plant genomes. Furthermore, it is not clear how the *cis*-regulatory elements within a single genome confer cell type-specific transcriptional activity – and thus cell type identity – during development. In the present study, we seek to build on previous work and to address some of these outstanding questions by analyzing chromatin accessibility across multiple, diverse plant species, and between two distinct cell types. From a methodological perspective, the DNase-seq procedure is relatively labor-intensive and requires a large number of starting nuclei for DNaseI treatment, which can be a major drawback for conducting cell type-specific profiling investigations. More recently, the Assay for Transposase-Accessible Chromatin with sequencing (ATAC-seq) was developed as an alternative approach (Buenrostro et al., 2013). ATAC-seq employs treatment of isolated nuclei with an engineered transposase that simultaneously cleaves DNA and inserts sequencing adapters, such that cleaved fragments originating from open chromatin can be converted into a high-throughput sequencing library by Polymerase Chain Reaction (PCR). Sequencing of the resulting library provides readout highly similar to that of DNase-seq, but ATAC-seq requires far fewer nuclei (Buenrostro et al., 2015). The relatively simple procedure for ATAC-seq and its low nuclei input, combined with its recent application in *Arabidopsis* and rice (Wilkins et al., 2016; Bajic et al., 2017; Lu et al., 2017), has made it widely useful for assaying plant DNA regulatory regions. In this study, we first optimized ATAC-seq for use with crude nuclei and nuclei isolated by INTACT (Isolation of Nuclei TAgged in specific Cell Types) affinity purification (Deal and Henikoff, 2010). We then applied this method to INTACT-purified root tip nuclei from *Arabidopsis thaliana*, *Medicago truncatula*, *Solanum lycopersicum* (tomato), and *Oryza sativa* (rice), as well as the root hair and non-hair epidermal cell types of *Arabidopsis*. The use of diverse plant species of both dicot and monocot lineages allowed us to assay regulatory structure over a broad range of evolutionary distances. Additionally, analysis of the *Arabidopsis* root hair and non-hair cell types allowed us to identify distinctions in chromatin accessibility that occurred during the differentiation of developmentally linked cell types from a common progenitor stem cell.

In our cross-species comparisons, we discovered that the majority of open chromatin sites in all four species exist outside of transcribed regions. The open sites also tended to cluster within several kilobases upstream of the transcription start sites despite the large differences in intergenic space between the four genomes. When orthologous genes were compared across species, we found that the number and location of open chromatin regions were highly variable, suggesting that regulatory elements are not statically positioned relative to target genes over evolutionary timescales. However, we found evidence that particular gene sets remain under control by common TFs across these species. For instance, we discovered a set of four TFs that appear to be integral for root tip transcriptional regulation of common gene sets in all species. These include HY5 and MYB77, which were previously shown to impact root development in *Arabidopsis* (Oyama et al., 1997; Shin et al., 2007).

When comparing the two *Arabidopsis* root epidermal cell types, we found that their open chromatin profiles are qualitatively very similar. However, many quantitative differences between cell types were identified, and these regions often contained binding motifs for TFs that were more highly expressed in one cell type than the other. Further analysis of several such cell type-enriched TFs led to the discovery of a hair cell transcriptional regulatory module driven by ABI5 and MYB33. These factors appear to co-regulate a number of additional hair cell-enriched TFs, including MYB44 and MYB77, which in turn regulate many downstream TF genes as well as other genes impacting hair-cell fate, physiology, secondary metabolism, and stress responses.

Overall, our work suggests that the *cis*-regulatory structure of these four plant genomes is strikingly similar, and that TF-target gene modules are also generally conserved across species. Furthermore, early differential expression of high-level TFs between the *Arabidopsis* hair and non-hair cells appears to drive a TF cascade that at least partially explains distinctions between hair and non-hair cell transcriptomes. Our data also highlight the utility of comparative chromatin profiling approaches and will be widely useful for hypothesis generation and testing.

## RESULTS AND DISCUSSION

### Application of ATAC-seq in *Arabidopsis* root tips

The Assay for Transposase-Accessible Chromatin (ATAC-seq) method was introduced in 2013 and has since been widely adopted in many systems (Buenrostro et al., 2013; Mo et al., 2015; Scharer et al., 2016; Lu et al., 2017). This technique utilizes a hyperactive Tn5 transposase that is pre-loaded with sequencing adapters as a probe for chromatin accessibility. When purified nuclei are treated with the transposase complex, the enzyme freely enters the nuclei and cleaves accessible DNA, both around nucleosomes and at nucleosome-depleted regions arising from the binding of transcription factors (TFs) to DNA. Upon cleavage of DNA, the transposon integrates sequencing adapters, fragmenting the DNA sample in the process. Regions of higher accessibility will be cleaved by the transposase more frequently and generate more fragments – and ultimately more reads, once the sample is sequenced. Conversely, less accessible regions will have fewer fragments and reads. After PCR-amplification of the raw DNA fragments, paired-end sequencing of the ATAC-seq library can reveal nucleosome-depleted regions where TFs are bound.

In this study, we set out to apply ATAC-seq to multiple plant species as well as different cell types from a single species. As such, we first established procedures for using the method with *Arabidopsis*, starting with root tip nuclei affinity-purified by INTACT (Isolation of Nuclei TAgged in specific Cell Types). We also established a protocol to use nuclei purified by detergent lysis of organelles followed by sucrose sedimentation, with the goal of broadening the application of ATAC-seq to non-transgenic starting tissue. We began with an *Arabidopsis* INTACT transgenic line constitutively expressing both the nuclear envelope targeting fusion protein (NTF) and biotin ligase (BirA) transgenes. Co-expression of these transgenes results in all the nuclei in the plant becoming biotinylated, and thus amenable to purification with streptavidin beads (Deal and Henikoff, 2010; Sullivan et al., 2014). Transgenic INTACT plants were grown on vertically oriented nutrient agar plates to facilitate root growth, and total nuclei were isolated from the 1 cm root tip region. These nuclei were further purified either by treatment with 1% (v/v) Triton X-100 and sedimentation through a sucrose cushion (‘Crude’ purification) or affinity-purified using streptavidin-coated magnetic beads (INTACT purification). In both cases 50,000 nuclei from each purification strategy were used as the input for ATAC-seq (Figure 1A). Overall, both Crude and INTACT-purified nuclei yielded very similar results (Figure 1B and C, Figure S1). One clear difference that emerged was the number of reads that map to organellar DNA between the nuclei preparation methods. While the total reads of Crude nuclei preparations mapped approximately 50% to organellar genomes and 50% to the nuclear genome, the total reads of INTACT-purified nuclei consistently mapped over 90% to the nuclear genome (Table 1). The issue of organellar genomes contaminating ATAC-seq reactions is a common one, resulting in a large percentage of organelle-derived reads that must be discarded before further analysis. This issue was also recently shown to be remedied by increasing the purity of nuclei prior to ATAC-seq by use of fluorescence-activated nuclei sorting (Lu et al., 2017). To compare between datasets for the Crude and INTACT preparation strategies, we analyzed the enrichment of ATAC-seq reads using Hotspot peak mapping software (John et al., 2011). Though designed for use with DNase-seq data, Hotspot can also be readily used with ATAC-seq data. The number of enriched regions found with this algorithm did not differ greatly between nuclei preparation types, nor did the SPOT score (a signal-specificity measurement representing the proportion of sequenced reads that fall into enriched regions) (Table 1). These results suggest that the datasets are generally comparable regardless of the nuclei purification method.

**Figure 1.**
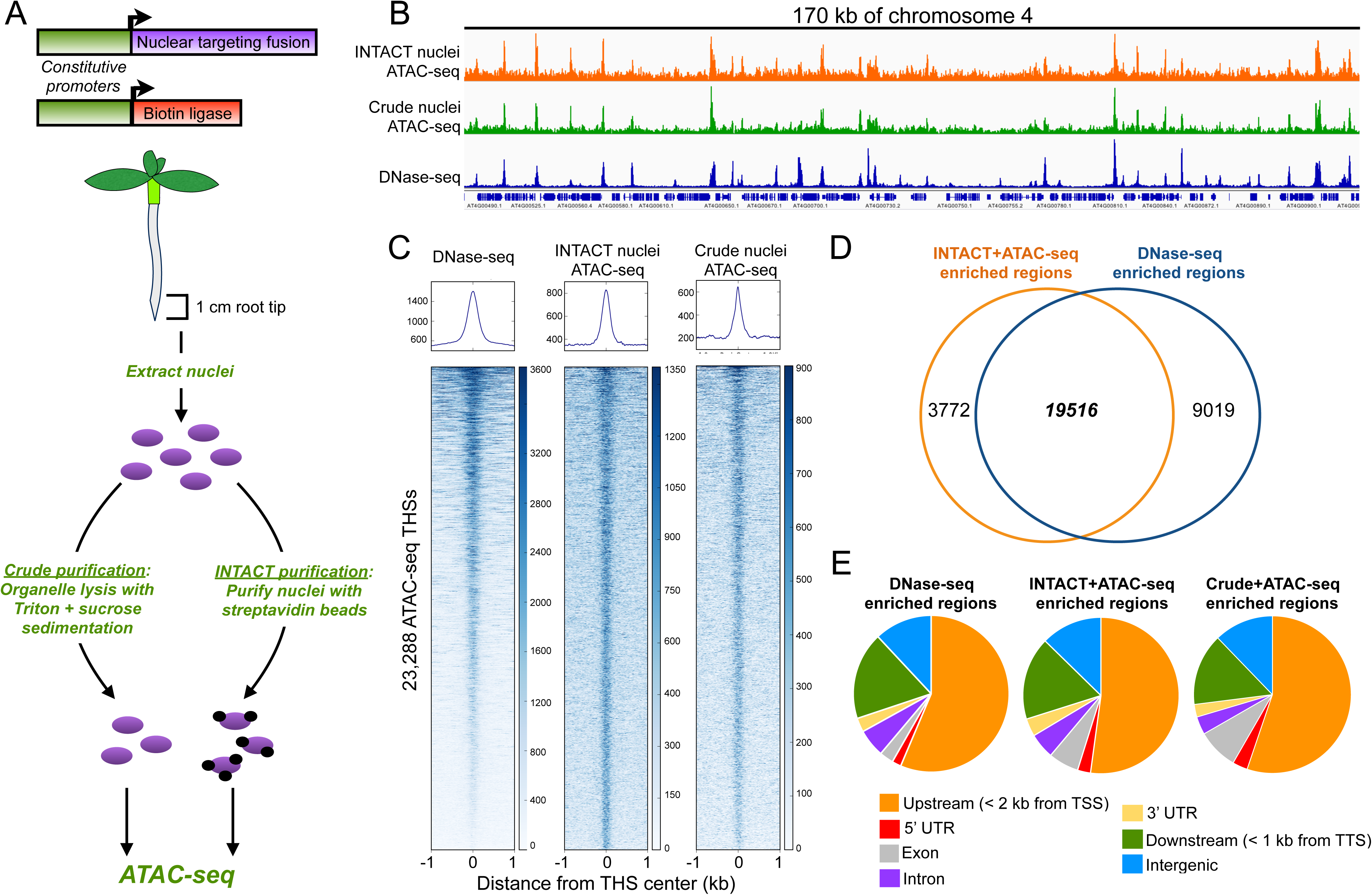
Application of ATAC-seq to *Arabidopsis* and comparison with DNase-seq data. **(A)** Schematic of the INTACT system and strategy for testing ATAC-seq on nuclei with different levels of purity. Upper panel shows the two transgenes used in the INTACT system: the nuclear targeting fusion (NTF) and biotin ligase. Driving expression of both transgenes using constitutive promoters generates biotinylated nuclei in all cell types. Below is a diagram of a constitutive INTACT transgenic plant, showing the 1 cm root tip section used for all nuclei purifications. Root tip nuclei were isolated from transgenic plants and either purified by detergent lysis of organelles followed by sucrose sedimentation (Crude) or purified using streptavidin beads (INTACT). In each case 50,000 purified nuclei were used as input for ATAC-seq. **(B)** Genome browser shot of ATAC-seq data along a 170 kb stretch of chromosome 4 from INTACT-purified and Crude nuclei, as well as DNase-seq data from whole root tissue. Gene models are displayed on the bottom track. **(C)** Average plots and heatmaps of DNase-seq and ATAC-seq signals at the 23,288 ATAC-seq transposase hypersensitive sites (THSs) in the INTACT-ATAC-seq dataset. The regions in the heatmaps are ranked from highest DNase-seq signal (top) to lowest (bottom) **(D)** Venn diagram showing the overlap of enriched regions identified in root tip INTACT-ATAC-seq and whole root DNase-seq datasets. **(E)** Genomic distributions of enriched regions identified in DNase-seq, INTACT-ATAC-seq, and Crude-ATAC-seq datasets.

**Table 1.** ATAC-seq reads from Crude and INTACT-purified *Arabidopsis* root tip nuclei. ATAC-seq was performed in biological triplicate for both Crude and INTACT-purified nuclei. For each replicate the table shows the percentage of reads mapping to organelle and nuclear genomes, the total number of enriched regions identified by the peak calling program Hotspot, as well as the SPOT score for each dataset. The SPOT score is a measure of specificity describing the proportion of reads that fall in enriched regions, with higher scores indicating higher specificity.

| Experiment | Plastid mapped reads (%) | Mitochondrial mapped reads (%) | Nuclear mapped reads (%) | Total nuclear mapped reads (x 10<sup>6</sup>) | Total Hotspot enriched regions called | SPOT score |
| --- | --- | --- | --- | --- | --- | --- |
| Crude 1 | 25.33 | 22.15 | 52.52 | 40.6 | 43,599 | 0.4339 |
| Crude 2 | 24.40 | 21.03 | 54.58 | 31.0 | 43,043 | 0.4086 |
| Crude 3 | 25.13 | 23.17 | 51.70 | 35.8 | 42,469 | 0.4471 |
| INTACT 1 | 4.62 | 2.44 | 92.94 | 34.6 | 36,463 | 0.4167 |
| INTACT 2 | 3.51 | 2.03 | 94.46 | 34.0 | 41,305 | 0.4004 |
| INTACT 3 | 2.81 | 1.61 | 95.57 | 89.7 | 55,857 | 0.4896 |

Visualization of the Crude- and INTACT-ATAC-seq datasets in a genome browser revealed that they were highly similar to one another and to DNase-seq data from whole root tissue (Figure 1B). Further evidence of similarity among these datasets was found by examining the normalized read count signal in all datasets (both ATAC-seq and DNase-seq) within the regions called as ‘enriched’ in the INTACT-ATAC-seq dataset. For this and all subsequent peak calling in this study, we used the *findpeaks* algorithm in the HOMER package (Heinz et al., 2010), which we found to be more versatile and user-friendly than Hotspot. Using this approach, we identified 23,288 enriched regions in our INTACT-ATAC-seq data. We refer to these peaks, or enriched regions, in the ATAC-seq data as transposase hypersensitive sites (THSs). We examined the signal at these regions in the whole root DNase-seq dataset and both Crude- and INTACT-ATAC-seq datasets using heatmaps and average plots. These analyses showed that THSs detected in INTACT-ATAC-seq tended to be enriched in both Crude-ATAC-seq and DNase-seq signal (Figure 1C). In addition, the majority of enriched regions (19,516 of 23,288) were found to overlap between the root-tip INTACT-ATAC-seq and the whole-root DNase-seq data (Figure 1D) and the signal intensity over DNase-seq or ATAC-seq enriched regions was highly correlated between the datasets (Figure S1). To examine the distribution of hypersensitive sites among datasets, we identified enriched regions in both types of ATAC-seq datasets and the DNase-seq dataset, and then mapped these regions to genomic features. We found that the distribution of open chromatin regions relative to gene features was nearly indistinguishable among the datasets (Figure 1E). In all cases, the majority of THSs (~75%) were outside of transcribed regions, with most falling within 2 kb upstream of a transcription start site (TSS) and within 1 kb downstream of a transcript termination site (TTS).

Overall, these results show that ATAC-seq can be performed effectively using either Crude or INTACT-purified nuclei, and that the data in either case are highly comparable to that of DNase-seq. While the use of crudely purified nuclei should be widely useful for assaying any tissue of choice without a need for transgenics, it comes with the drawback that ~50% of the obtained reads will be from organellar DNA. The use of INTACT-purified nuclei greatly increases the cost efficiency of the procedure and can also provide access to specific cell types, but requires pre-established transgenic lines.

### Comparison of root tip open chromatin profiles among four species

Having established an efficient procedure for using ATAC-seq on INTACT affinity-purified nuclei, we used this tool to compare the open chromatin landscapes among four different plant species. In addition to the *Arabidopsis* INTACT line described above, we also generated constitutive INTACT transgenic plants of *Medicago truncatula* (*Medicago*), *Oryza sativa* (rice), and *Solanum lycopersicum* (tomato). Seedlings of each species were grown on vertically oriented nutrient plates for one week after radicle emergence, and nuclei from the 1 cm root tip regions of each seedling were isolated and purified with streptavidin beads. ATAC-seq was performed in at least two biological replicates for each species, starting with 50,000 purified nuclei in each case. Visualization of the mapped reads across each genome showed notable consistencies in the data for all four species. In all cases, the reads localize to discrete peaks that are distributed across the genome, as expected (Figure 2A). Examination of a syntenic region found in all four genomes suggested at least some degree of consistency in the patterns of transposase accessibility around orthologous genes (Figure 2A).

**Figure 2.**
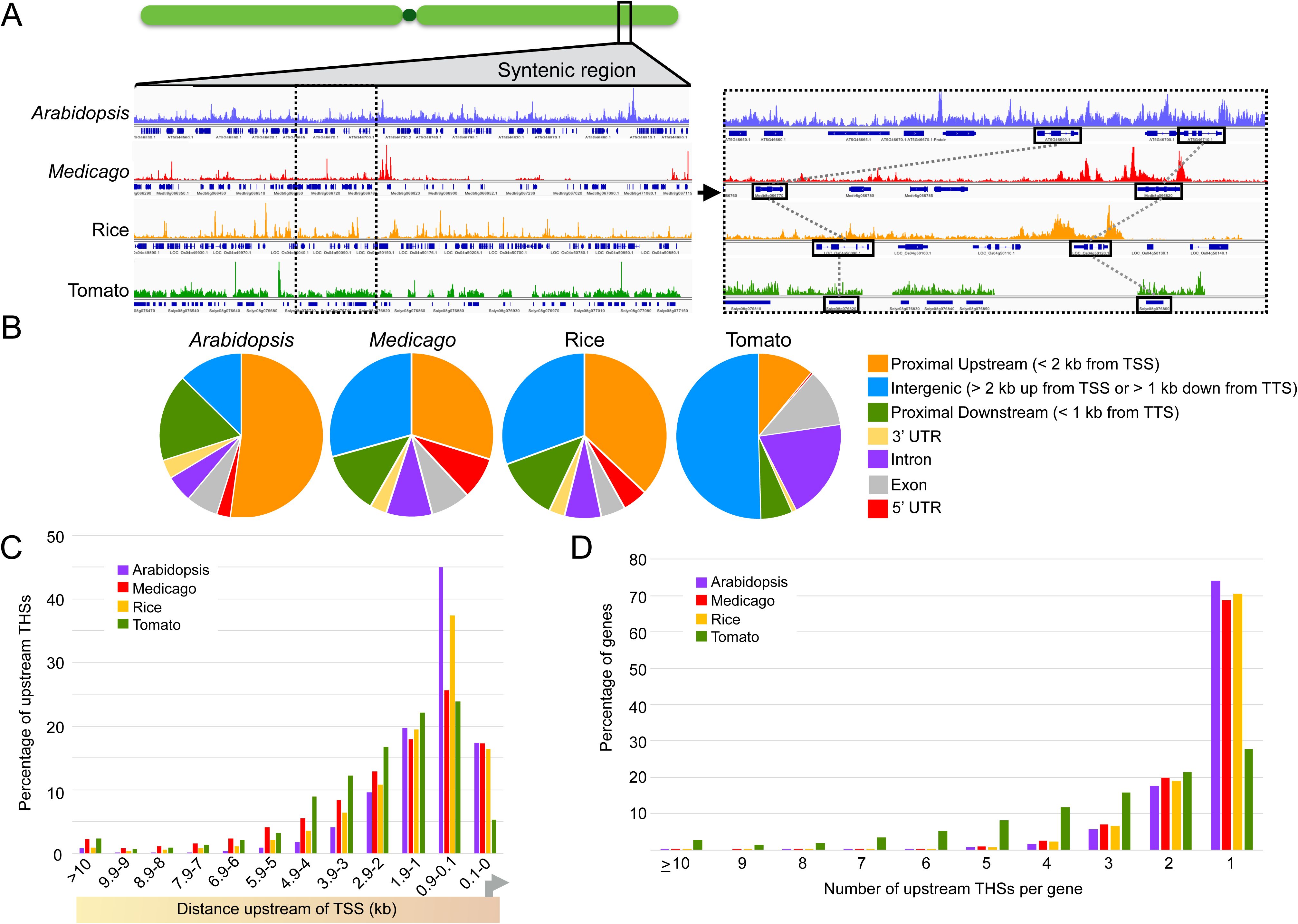
ATAC-seq profiling of *Arabidopsis*, *Medicago*, tomato, and rice. **(A)** Comparison of ATAC-seq data along syntenic regions across the species. The left panel shows a genome browser shot of ATAC-seq data across a syntenic region of all four genomes. ATAC-seq data tracks are shown above the corresponding gene track for each species. The right panel is an enlargement of the region surrounded by a dotted box in the left panel. Orthologous genes are surrounded by black boxes connected by dotted lines between species. Note the apparent similarity in transposase hypersensitivity upstream and downstream of the rightmost orthologs. **(B)** Distribution of ATAC-seq transposase hypersensitive sites (THSs) relative to genomic features in each species. **(C)** Distribution of upstream THSs relative to genes in each species. THSs are binned by distance upstream of the transcription start site (TSS). The number of peaks in each bin is expressed as a percentage of the total upstream THS number in that species. **(D)** Number of upstream THSs per gene in each species. Graph shows the percentage of all genes with a given number of upstream THSs.

To specifically identify regions of each genome that were enriched in ATAC-seq signal (THSs), we used the HOMER *findpeaks* function on each biological replicate experiment. For further analysis, we retained only THS regions that were found in at least two biological replicates of ATAC-seq in each species. These reproducible THSs were then mapped to genomic features in each species in order to examine their distributions. As seen previously for *Arabidopsis*, the majority of THSs (~70–80%) were found outside of transcribed regions in all four species (Figure 2B). For this analysis, we classified these extragenic THSs (THSs found anywhere outside of transcribed regions) as proximal upstream (< 2 kb upstream of the transcription start site, or TSS), proximal downstream (< 1 kb downstream of the transcript termination site, or TTS) or intergenic (> 2 kb upstream from a TSS or > 1 kb downstream from a TTS). The proportion of THSs in the proximal upstream and intergenic regions varied greatly with genome size, and thus the amount of intergenic space in the genome. For example, a full 52% of THSs in *Arabidopsis –* the organism with the smallest genome (~120 Mb) and highest gene density of the four species – were in the proximal upstream region. This percentage drops as genome size and intergenic space increase, with 37% of the THSs in the proximal upstream region in the rice genome (~400 Mb), 30% in the *Medicago* genome (~480 Mb), and a mere 11% in the tomato genome (~820 Mb). The percentage of total THSs in the proximal downstream region followed a similar pattern, marking 17% of the THSs in *Arabidopsis*, 12% in rice and *Medicago*, and 6% in tomato. Finally, the proportion of THSs classified as intergenic followed the inverse trend as expected, with 12% of the THSs in intergenic regions for *Arabidopsis*, 30% for rice and *Medicago*, and 50% for tomato (Figure 2B).

Thus, while the overall proportion of extragenic THSs is similar among species, the distance of these sites from genes tends to increase with genome size, which is roughly proportional to the average distance between genes.

Since the majority of THSs were found upstream of the nearest gene for each species, we next classified the regions based on their distance from the nearest TSS. We binned THSs in each genome into twelve distance categories, starting with those > 10 kb upstream of the TSS, then into eleven bins of 999 bp moving in toward the TSS, and finally a TSS-proximal bin of 100-0 bp upstream of the TSS (Figure 2C). Starting with this TSS-proximal bin, we find that ~17% of the upstream THSs in *Arabidopsis*, *Medicago*, and rice are within 100 bp of the TSS, whereas 2.7% of the upstream THSs in tomato are within 100 bp of the TSS. Moving away from the TSS, we find that 91% of the total upstream THSs fall within 2.9 kb of the TSS in *Arabidopsis*, while this number decreases with genome size, with 84% for rice, 73% for *Medicago*, and 65% for tomato. In the distance bin spanning 9.9 kb to 3 kb upstream, we find 7% of the total upstream THSs in *Arabidopsis*, 15% in rice, 23% in *Medicago*, and 32% in tomato. Finally, the THSs that are more than 10 kb away from the TSS accounts for 0.8% of the total upstream THSs in *Arabidopsis*, 0.9% in rice, 2.3% in *Medicago*, and 3.3% in tomato. Overall, it is clear that in all species the majority of THSs are within 3 kb upstream of a TSS, suggesting that most *cis*-regulatory elements in these genomes are likely to be proximal to the core promoter. In the species with the largest genomes and intergenic distances (*Medicago* and tomato), THSs tend to be spread over a somewhat wider range upstream of the TSS. However, even in these cases, only a few hundred THSs in total are more than 10 kb away from the nearest gene. It is worth noting that the distribution of THSs in *Medicago* is more similar to that of tomato than rice, despite the genome size being more similar to rice. This suggests that THSs tend to be further away from TSSs in *Medicago* than would be expected based on genome size alone.

As most THSs fall near genes, we next investigated from the opposite perspective – for any given gene, how many THSs were associated with it? In this regard, we find that the *Arabidopsis*, *Medicago*, and rice genomes are highly similar (Figure 2D). In all three genomes, of the subset of genes that have *any* upstream THSs, ~70% of these genes have a single site, ~20% have two sites, 5–7% have three sites, and 2–3% have four or more THSs. By contrast, the tomato genome has a different trend. Of the subset of tomato genes with *any* upstream THS, only 27% of the genes have a single site, and this proportion gradually decreases with increasing THS number, with 2.7% of the tomato genes in this subset having 10 or more THSs.

Overall, we have found that THSs have similar size and genomic distribution characteristics across all four species (Table S1). The majority of THSs in all species are found outside of genes, mainly upstream of the TSS, and these sites tend to cluster within 3 kb of the TSS. Furthermore, most genes with an upstream THS in *Arabidopsis*, *Medicago*, and rice have only 1–2 THSs, whereas tomato genes tend to have a larger number of upstream THSs. Whether this increase in upstream THSs in tomato is reflective of an increase in the number of regulatory elements per gene based on clade-specific alterations in gene regulation, DNA copy number changes, or simply the greater abundance of transposons and other repeat elements is not entirely clear. Compared to the other species, tomato THSs are much more abundant and tend to be smaller in size than those of the other species, and the tomato ATAC-seq data generally appear to have a lower signal-to-noise ratio (Table S8, Figure 2A). While it is unclear why the data from tomato are distinct in these ways, it is clear that tomato THSs occupy mostly genic regions of the genome, as expected, and are highly reproducible between biological replicate experiments (Figure S2).

Collectively, these results suggest that there is a relatively small number of regulatory elements per gene in plants. These elements tend to be focused near the promoter rather than at more distal sites as has been observed in animal, particularly mammalian, genomes (Stadhouders et al., 2012). The assumptions implicit in this argument are that open chromatin sites near a TSS reflect regulatory elements that regulate that TSS and not a more distant one, and that upstream elements contribute the majority of regulatory effects. These assumptions appear to be generally validated by many reporter assays showing that an upstream fragment of several kilobases is frequently sufficient to recapitulate native transcription patterns (Medford et al., 1991; Masucci et al., 1996; Ruzicka et al., 2007; Tittarelli et al., 2009; Li et al., 2012), as well as our observation that upstream THSs are the most abundant class of open chromatin sites.

### Open chromatin features are not directly conserved among orthologous genes

Given that many of the properties of open chromatin regions were shared among *Arabidopsis*, *Medicago*, rice, and tomato, we next asked whether the numbers and locations of THSs – and thus putative regulatory elements – were conserved among orthologous genes across species. For these analyses, we identified 373 syntenic orthologs (Table S2) that were found in all four genomes and asked whether members of each ortholog set harbored a similar number of open chromatin regions across the species. Again, using root tip THSs present in at least two biological replicates for each species, we counted the number of THSs within 5 kb upstream of the TSS for each ortholog in each species. We then examined these data for similarities and differences in upstream THS number (Figure 3A). While no clear trend of strong conservation in the number of upstream THSs emerged from this analysis, there was a small subset of orthologs that did have upstream THSs in similar numbers across species. However, this was a very small proportion of the total. As seen in earlier analyses, tomato genes tended to have a larger number of upstream THSs compared to the other species, and most of the 373 orthologs in tomato did have at least one upstream THS. This was not the case in the other three species, where many of the orthologs had no detectable upstream THSs within 5 kb of the TSS. Among the four species, *Arabidopsis* and *Medicago* showed the greatest similarity in upstream THS number, but even in this case the similarity was minimal despite the relatively closer phylogenetic relationship between these two organisms.

**Figure 3.**
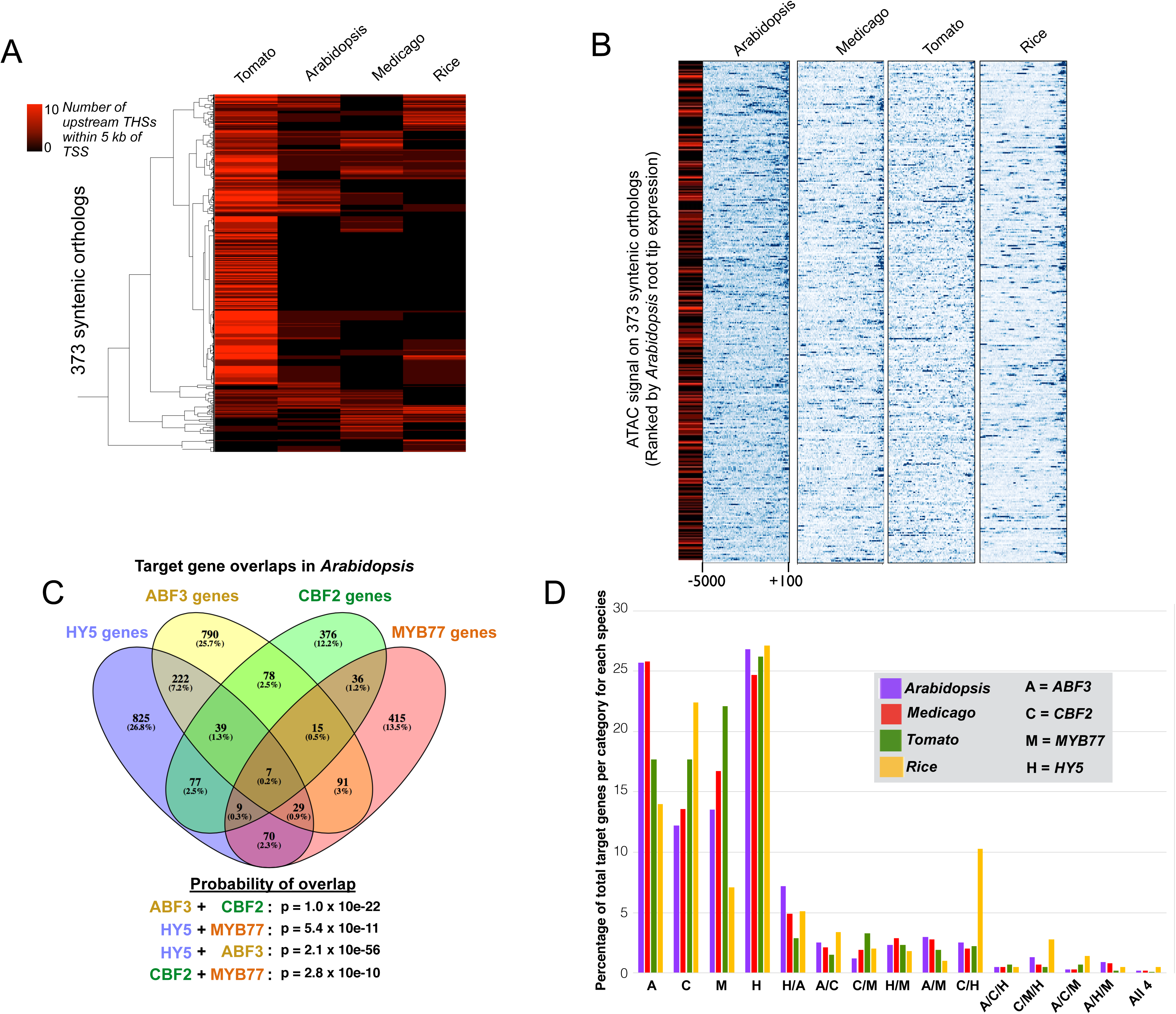
Characterization of open chromatin regions and regulatory elements in *Arabidopsis*, *Medicago*, tomato, and rice. **(A)** Heatmap showing the number of upstream THSs at each of 373 syntenic orthologs in each species. Each row of the heatmap represents a syntenic ortholog, and the number of THSs within 5 kb upstream of the TSS is indicated with a black-to-red color scale for each ortholog in each species. Hierarchical clustering was performed on orthologs using uncentered correlation and average linkage. **(B)** Normalized ATAC-seq signals upstream of orthologous genes. Each row of the heatmaps represents the upstream region of one of the 373 syntenic orthologs in each species. ATAC-seq signal is shown across each ortholog from +100 to −5000 bp relative to the TSS, where blue is high signal and white is no signal. Heatmaps are ordered by transcript level of each *Arabidopsis* ortholog in the root tip, from highest (top) to lowest (bottom). The leftmost heatmap in black-to-red scale indicates the number of upstream THSs from −100 to −5000 bp associated with each of the *Arabidopsis* orthologs, on the same scale as in (A). **(C)** Overlap of predicted target genes for HY5, ABF3, CBF2, and MYB77 in the *Arabidopsis* root tip. Predicted binding sites for each factor are those THSs that also contain a significant motif occurrence for that factor. Venn diagram shows the numbers of genes with predicted binding sites for each factor alone and in combination with other factors. Significance of target gene set overlap between each TF pair was calculated using a hypergeometric test with a population including all *Arabidopsis* genes reproducibly associated with an ATAC-seq peak in the root tip (13,714 total genes). For each overlap, we considered all genes co-targeted by the two factors. **(D**) Conveying data similar to that in (C), the clustered bar graph shows the percentage of total target genes that fall into a given regulatory category (targeted by a single TF or combination of TFs) in each species.

We next examined the distribution of open chromatin regions across the upstream regions of these 373 orthologous genes relative to their expression level in *Arabidopsis*, reasoning that there could be patterns of open chromatin similarity based on THS positions, rather than numbers. For this analysis, we examined the normalized ATAC-seq signal across the upstream region of all 373 orthologous genes, from −5000 bp to +100 bp relative to the TSS of each gene (Figure 3B). Orthologs were then ranked within the heatmap based on the transcript level of each *Arabidopsis* ortholog in the root tip (Li et al., 2016), from highest to lowest expression. For each *Arabidopsis* ortholog we also included the upstream THS number to ascertain how this feature might correlate with transcript level for *Arabidopsis*. While there was some consistency among species in that open chromatin often overlapped with the TSS, we did not observe any clear pattern in transposase hypersensitivity within the upstream regions of these orthologs. K-means clustering of the heatmaps similarly did not reveal evidence for conservation of open chromatin patterns among orthologs (Figure S3A). An important caveat to this analysis is that many of these syntenic orthologs may not be functional homologs, or ‘expressologs’ (Patel et al., 2012), due to subfunctionalization within gene families. As such, we identified a smaller group (52) of expressologs on which to perform a similar test (Table S3). While these expressolog genes have both maximally high protein level similarity and expression pattern similarity, including expression in the root, there was also no clear correspondence in upstream THS number among them (Figure S3B).

There does not appear to be strong conservation in the number and location of open chromatin sites at orthologous genes across species. Assuming that these genes are still under control of common TFs, this suggests that regulatory elements could be free to migrate, and perhaps split or fuse, while retaining the regulatory parameters of the target gene in question.

One interesting finding from these analyses was that the pattern of upstream THS number does not correlate with expression level, at least for *Arabidopsis* (Figure 3B). Thus, THSs must not simply represent activating events upstream of the TSS but may also represent binding of repressive factors. Further, we found no correlation between upstream THS number and expression entropy among all genes in the *Arabidopsis* genome, suggesting a more complex relationship between regulatory element distribution and target gene transcription (Figure S3C).

### Evidence for co-regulation of common gene sets by multiple TFs across species

While there does not appear to be a consistent pattern in the number or placement of open chromatin regions around orthologs or expressologs, we wanted to examine whether it would be possible to find common regulators of specific gene sets among species using a deeper level of analysis. To do this, we first searched for common TF motifs in root tip THSs across the four species. Using the THSs that were found in at least two replicates for each species, we employed the MEME-ChIP motif analysis package (Machanick and Bailey, 2011; Ma et al., 2014) to identify overrepresented motifs of known TFs. We discovered 30 motifs that were both overrepresented and common among all species (Table S4). We narrowed our list of candidate TFs by considering a variety of factors, including the expression of each TF in the root tip, any known mutant root phenotypes involving those TFs, and whether genome-wide binding information was available for each candidate in *Arabidopsis*. Ultimately, we selected 4 TFs for further analysis: ELONGATED HYPOCOTYL 5 (HY5), ABSCISIC ACID RESPONSIVE ELEMENTS-BINDING FACTOR 3 (ABF3), C-REPEAT/DRE BINDING FACTOR 2 (CBF2), and MYB DOMAIN PROTEIN 77 (MYB77). It is worth noting that among these factors, both HY5 and MYB77 had been previously implicated in root development (Oyama et al., 1997; Zhao et al., 2014). Like HY5 and MYB77, CBF2 and ABF3 have been implicated in stress responses as well as abscisic acid (ABA) signaling (Kang et al., 2002; Knight et al., 2004). Furthermore, overexpression of ABF3 leads to increased tolerance to multiple abiotic stresses in *Arabidopsis*, rice, cotton, and alfalfa (Oh et al., 2005; Abdeen et al., 2010; Wang et al., 2016; Kerr et al., 2017). Given this evidence, we decided to focus on these factors for further study.

We first sought to define the target genes for each of these four TFs in *Arabidopsis* by combining our chromatin accessibility data with published genome-wide binding data for each factor in *Arabidopsis* (Table 2). Because an accessible chromatin region (a THS) represents the displacement of nucleosomes by a DNA-binding protein, we reasoned that our THS profiles for a given tissue would represent virtually all possible protein binding sites in the epigenomes of root tip cells. Similarly, by using *in vitro* genomic binding data (DAP-seq) (O’Malley et al., 2016b) or ChIP-seq data from a highly heterogeneous tissue, we could identify the spectrum of possible binding sites for that TF, such that the intersection of these datasets would represent the binding sites for that TF in the sample of interest. While there are caveats to this approach, we reasoned that it was more likely to generate false negatives than false positives and would give us a set of high confidence target genes to analyze for each TF. In this regard, ChIP-seq data may be more robust because they represent *in vivo* binding, while DAP-seq is an *in vitro* assay and may not capture binding sites that depend on chromatin properties or interactions with other TFs. On the other hand, ChIP-seq data are inherently limited by the cell types present in the sample used.

**Table 2.** Transcription factor motifs significantly enriched in transposase hypersensitive sites (THSs) in all four species. THSs found in at least two replicates for each species were analyzed for overrepresented TF motifs. Four of the thirty TFs that were significantly enriched in THSs of all four species are shown in the table. Significant occurrences of each TF motif were identified across the *Arabidopsis* genome, and the percentage of these motif occurrences that fall within known binding sites for that factor (based on published ChIP-seq or DAP-seq datasets) are indicated in Column 4. The final column indicates the percentage of *Arabidopsis* root tip THSs that contain a motif for each factor and also overlap with a known binding site for the factor. These are considered high-confidence binding sites (Figure S4).

| Transcription Factor | Family | Average expression in *Arabidopsis* root tip (RPKM) | Percentage of motif occurrences in the *Arabidopsis* genome that overlap with a known binding site | Percentage of motif-containing THSs in *Arabidopsis* that overlap with a known binding site (high-confidence binding sites) |
| --- | --- | --- | --- | --- |
| AT5G11260 (HY5) | bZIP | 37 | 30.3%<br>(7,156/23,541) | 61.2%<br>(810/1,323) |
| AT4G34000 (ABF3) | bZIP | 1 | 53.3%<br>(3,821/7,164) | 97.1%<br>(1,279/1,316) |
| AT4G25470 (CBF2) | AP2 | 16 | 64.2%<br>(13,144/20,457) | 89.1%<br>(582/653) |
| AT3G50060 (MYB77) | MYB/SANT | 6 | 52.5%<br>(9,147/17,402) | 74.6%<br>(506/678) |

We first tested this approach in *Arabidopsis* with each of the four TFs of interest. Using THSs from the *Arabidopsis* root tip that were found in at least two biological replicates, we used the motif-identification tool FIMO (Grant et al., 2011) to identify THSs that contained a significant occurrence of the TF motif of interest. The THSs that contained a significant motif match were considered *predicted binding sites*. We then identified predicted binding sites that also overlapped with a known binding site for that TF (a DAP-seq or ChIP-seq peak), and these were considered *high confidence binding sites* for that TF in the root tip (Figure S4). The predicted binding sites (motif-containing THSs) were themselves very good predictors of the true binding sites for these four TFs (Table 2). For example, of the 1,316 *Arabidopsis* root tip THSs with an occurrence of the ABF3 motif (Mathelier et al., 2014), 1,279 (97%) overlapped with an ABF3 ChIP-seq peak from whole 2-day-old seedlings (Song et al., 2016). Similarly, 89% of predicted CBF2 binding sites (Weirauch et al., 2014a) overlapped with a CBF2 DAP-seq peak (O'Malley et al., 2016a), 74% of predicted MYB77 binding sites (Weirauch et al., 2014a) overlapped with a MYB77 DAP-seq peak (O'Malley et al., 2016a), and 61% of predicted HY5 binding sites (Mathelier et al., 2014) overlapped with a HY5 DAP-seq peak (O'Malley et al., 2016a). In each case, the high confidence binding sites (motif-containing THSs that overlap with a ChIP- or DAP-seq peak) were assigned to their nearest TSS in order to identify the putative target genes for each TF (Figure S4).

With these lists of target genes for each TF in the *Arabidopsis* root tip, we looked for gene sets that were regulated by more than one factor, as means of identifying co-regulatory associations between these four TFs. We found extensive co-targeting among these four TFs, with gene sets being targeted by one, two, three, or all four of these TFs to a degree that was far higher than what would be expected by chance (Figure 3C). For example, of the 1,271 ABF3 target genes, 297 (23%) are also targeted by HY5 (hypergeometric *p* = 2.1 × 10^−56^). Among these 297 genes, 46 are targeted by ABF3, HY5, and CBF2, and seven are targeted by all four TFs. We also asked where the binding sites driving this pattern were located relative to the target genes. To do this we considered only binding sites within the 5 kb upstream region of a TSS, and repeated the target gene assignment and analysis of target gene overlaps between TFs. This subsetting reduced the total number of target genes for each factor by ~20%, but did not substantially alter the percentages of target gene overlap among the four TFs (Figure S5A). These results collectively suggest that these four TFs have important roles in root tip gene regulation both individually and in combination, and that the majority of their binding sites (~80%) fall within the 5 kb region upstream of the TSS for target genes. In addition, we find that the binding sites for multiple TFs often occur in the same THS (Figure S5B).

We next sought to examine the target genes and proportions of target gene overlaps between the four species to address the conservation of co-regulatory relationships among these four TFs. Given that no TF binding data is available for the other three species and knowing that the majority of our predicted binding sites in *Arabidopsis* corresponded to known binding sites (Table 2; 61–97%), we opted to also use the predicted binding sites for each of the four TFs in *Medicago*, tomato, and rice, with the knowledge that these sets may contain some false positives. For these analyses, we used the *Arabidopsis* TF motifs – since these have not been directly defined for the other species – with the caveat that the DNA binding specificity of these factors may not be identical among species.

We again used FIMO to identify significant occurrences of each TF motif within the root tip THSs found in at least two biological replicates for each of our four species. We then mapped the predicted binding sites of each TF to the nearest TSS to define target genes for each TF in each species (Table S5). We then analyzed the overlap of TFs at target genes in each species using 4-way Venn diagrams, similar to Figure 3C. To compare regulatory associations across species, we considered each of the 15 categories in every species-specific 4-way Venn diagram as a regulatory category. For example, one regulatory category consists of the genes targeted only by ABF3 alone, another would be those targeted only by HY5 and ABF3 at the exclusion of the other two TFs, and so on. For each regulatory category in each species, we calculated the percentage of the total target genes in that category (number of genes in the regulatory category/total number of genes targeted by *any* of the 4 TFs), and then compared these percentages between species (Figure 3D). We found remarkably consistent proportions of the target genes in nearly all regulatory categories across all four species. However, notable deviations from this consistency among species were seen in the proportion of rice genes targeted by MYB77 alone and rice genes targeted CBF2 and HY5 together. In most cases, the proportions of target genes in different regulatory categories were most similar between *Arabidopsis* and *Medicago*, and these were generally more similar to tomato than to rice, consistent with the evolutionary distances between the species (Vanneste et al., 2014). Commonly overrepresented Gene Ontology (GO) terms among gene sets in particular regulatory categories across species further support the notion of regulatory conservation (Figure S5C), although these analyses are limited by the depth of GO annotation in some of these species.

These findings suggest that while neither syntenic orthologous gene sets nor expressolog gene sets tend to share open chromatin patterns, the genes under control of specific TFs or specific combinations of TFs appear to be relatively stable over evolutionary time, at least for the four TFs we examined. One simple explanation for this phenomenon is that the locations of transcriptional regulatory elements are somewhat malleable over time as long as proper transcriptional control is maintained. In this model, these elements would be free to relocate in either direction, and potentially even merge or split. This would maintain proper control over the target gene, but give each ortholog or expressolog a unique chromatin accessibility profile depending on the exact morphology and distribution of the functionally conserved regulatory elements. This idea of modularity is consistent with previous observations that the *Drosophila* even-skipped stripe 2 enhancer can be rearranged and still retain functionality (Ludwig et al., 2000; Ludwig et al., 2005). The results also shed light on the interconnectedness of specific TFs in root tip cells and indicate durability of these co-regulatory relationships over time. They also generate readily testable hypotheses regarding the how HY5, ABF3, MYB77, and CBF2 operate during root development. For example, given that HY5 appears to regulate over 1,000 genes in the *Arabidopsis* root tip (Figure 3C), and that hundreds of these are annotated with GO terms including *biological regulation* and *response to stimulus*, we predict that *hy5* mutants would have defects in root tip morphology and growth. Indeed, HY5 was previously shown to be involved in the regulation of lateral root growth initiation and gravitropism (Oyama et al., 1997), and we observe that the primary root tips in *hy5* mutants also frequently show a bulging and malformed appearance, as well as severe gravitropism defects (Figure S6).

### Commonalities and distinctions in the open chromatin landscapes of *Arabidopsis* root epidermal cell types

Having examined questions of regulatory conservation between species, we then explored regulatory elements and TFs relationships between cell types within a single species. In this case, we chose to focus on the root epidermal hair and non-hair cell types in *Arabidopsis*. Since these two cell types are derived from a common progenitor, they are prime candidates to offer insight into the epigenomic alterations that occur during – and likely drive – cell differentiation. Specifically, we investigated to what extent the open chromatin landscapes would differ between cell types and whether differences in THSs could pinpoint the sites of differential transcriptional regulation. Furthermore, we wanted to understand whether we could use this information to examine the TF-to-TF regulatory connections that underlie the transcriptomic and physiological differences between these cell types.

We used two previously described INTACT transgenic lines as starting material for these experiments: one having biotin-labeled nuclei exclusively in the root hair (H) cells, and another with labeled nuclei only in the root epidermal non-hair (NH) cells (Deal and Henikoff, 2010). Nuclei were purified from each fully differentiated cell type by INTACT, and 50,000 nuclei of each type were subjected to ATAC-seq. Visualization of these cell type-specific datasets in a genome browser, along with the *Arabidopsis* whole 1 cm root tip ATAC-seq data, showed a high overall degree of similarity among the three datasets (Figure 4A). Comparison of the ATAC-seq signal intensity at common THS regions genome-wide revealed that these two cell types have open chromatin patterns that are highly similar to one another, but distinct from that of the whole root tip (Figure S7).

**Figure 4.**
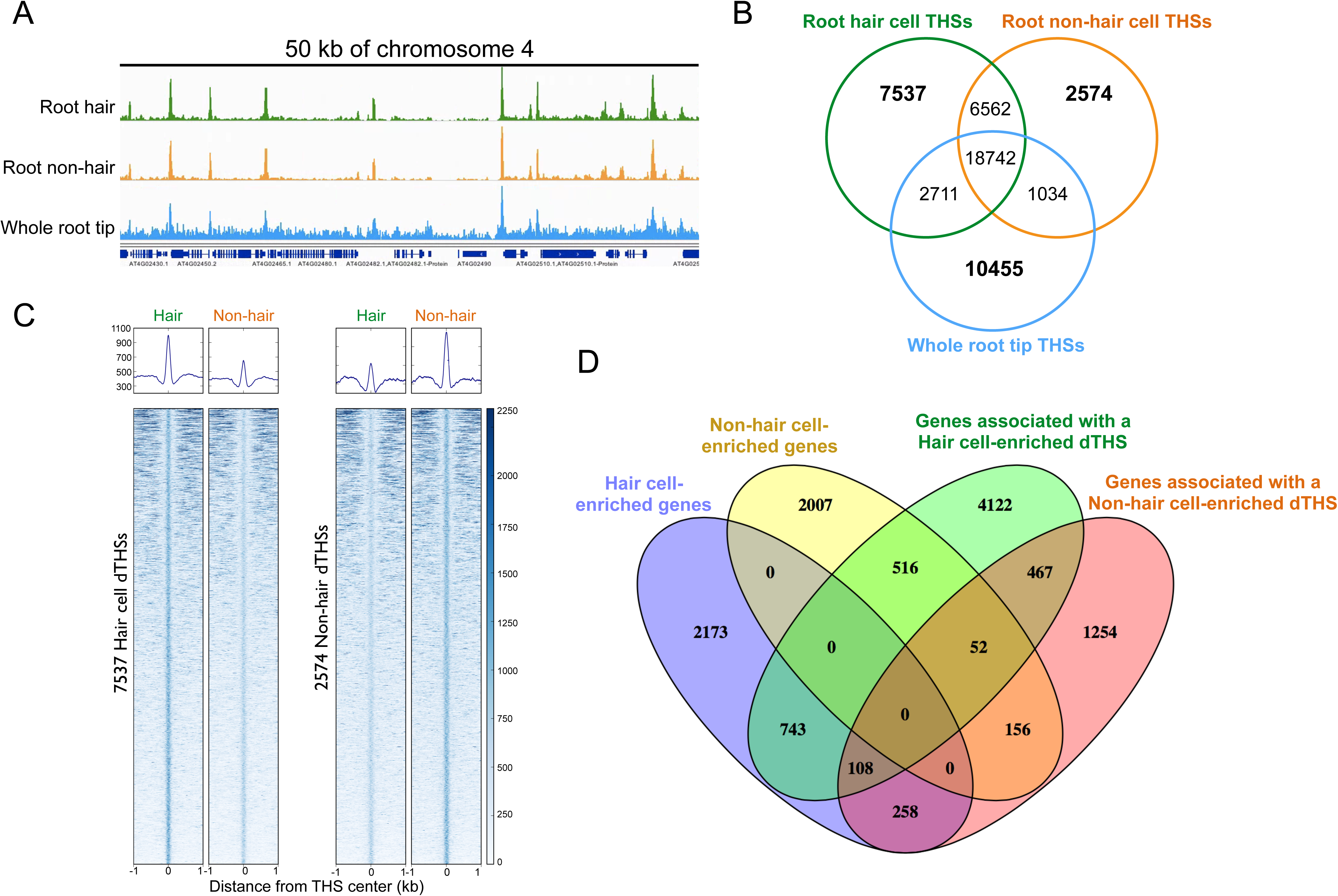
Characterization of open chromatin regions in the *Arabidopsis* root hair and non-hair cell types. **(A)** Genome browser shot of ATAC-seq data from root hair cell, non-hair cell, and whole root tip representing 50 kb of Chromosome 4. **(B)** Overlap of THSs found in two biological replicates of each cell type or tissue. Numbers in bold indicate THSs that are only found in a given cell type or tissue (differential THSs, or dTHSs). **(C)** Average plots and heatmaps showing normalized ATAC-seq signals over 7,537 root hair cell dTHSs (left panels) and 2,574 non-hair cell-enriched dTHSs (right panels). Heatmaps are ranked in decreasing order of total ATAC-seq signal in the hair cell panel in each comparison. Data from one biological replicate is shown here and both replicate experiments showed very similar results. **(D)** Venn diagram of overlaps between cell type-enriched gene sets and genes associated with cell type-enriched dTHSs. Transcriptome data from hair (purple) and non-hair cells (yellow) are from Li et al. (2016) *Developmental Cell*. Genes were considered cell type-enriched if they had a 2-fold or higher difference between cell types and a read count of 5 RPKM or greater in the cell type with higher expression.

To identify regions of differential accessibility between the cell types and the whole root tip, we considered THS regions that were found in at least two biological replicates of each cell type or tissue. The total number of these reproducible THSs was 32,942 in the whole root tip, 35,552 for the H cells, and 28,912 for the NH cells. The majority of these sites (18,742) were common (overlapping) in all three sample types (Figure 4B) and thus likely represent regulatory sites that are utilized in multiple *Arabidopsis* root cell types. We also found 6,562 THSs that were common to both root epidermal cell types but were not found in the whole root tip, suggesting that these may represent epidermal-specific regulatory elements. In a search for unique THSs in each of the three sample types (those not overlapping with a THS in any other sample), we found 10,455 THSs that were unique to the whole root tip, 7,537 unique to the H cells, and 2,574 that were unique to the NH cells. We refer to these regions as differential THSs (dTHSs). The dTHSs identified only in the H or NH cell type were of further interest because they may represent regulatory elements that drive the transcriptomic differences between these two epidermal cell types.

To examine the extent of chromatin accessibility differences at these dTHSs, we visualized the accessibility signals from each cell type at both H cell dTHSs and NH cell dTHSs. First, using the 7,537 regions identified as H cell dTHSs, we used heatmaps and average plots to examine the normalized ATAC-seq read count across these regions in each cell type (Figure 4C, left panel). We then repeated this analysis using the 2,574 NH cell dTHSs (Figure 4C, right panel). In each case, it was clear that the regions we identified as dTHSs showed significant differences in chromatin accessibility between the two cell types. However, the differences in chromatin accessibility between cell types were quantitative (varying intensity) rather than qualitative (all-or-nothing). This indicates that, at large, the dTHSs represent sites that are highly accessible in one cell type and less so in the other, rather than being strictly present in one and absent in the other. Therefore, we refer to these sites from this point on as cell type-enriched dTHSs to convey the notion of quantitative differences between cell types.

To identify the genes that might be impacted by cell type-enriched dTHSs, we mapped each dTHS to its nearest TSS and considered that to be the target gene. We found that the 7,537 H-enriched dTHSs mapped to 6,008 genes, while the 2,574 NH-enriched dTHSs mapped to 2,295 genes. Thus, the majority of genes that are associated with a dTHS are only associated with one such site. This is consistent with our previous findings that most *Arabidopsis* genes are associated with a single upstream THS (Figure 2D).

We then asked how the set of genes associated with dTHSs overlapped with those whose transcripts that show differential abundance between the two cell types. Using data from a recent comprehensive RNA-seq analysis of flow sorted *Arabidopsis* root cell types (Li et al., 2016a), we identified sets of transcripts that were more highly expressed in H versus NH cell types. To be considered a *cell type-enriched gene*, we required a gene to have a transcript level with two-fold or greater difference in abundance between H and NH cell types, as well as at least five reads per kilobase per million mapped reads (RPKM) in the cell type with a higher transcript level. Using this relatively conservative approach, we derived a list of 3,282 H cell-enriched genes and 2,731 NH cell-enriched genes. We then asked whether the genes associated with cell type-enriched dTHSs were also cell type-enriched genes (Figure 4D). Of the 3,282 H cell-enriched genes, 743 were associated with a H cell-enriched dTHS, 258 were associated with a NH cell-enriched dTHS, and 108 genes were associated with a dTHS in both cell types. Among the 2,731 NH cell-enriched genes, 156 were associated with a NH cell-enriched dTHS, 516 were associated with a H cell-enriched dTHS, and 52 genes showed dTHSs in both cell types. These results suggest that cell type-enriched expression of a gene is frequently associated with a dTHS in the cell type where the gene is highly expressed, but is also often associated with a dTHS in the cell type where that gene is repressed. This highlights the importance of transcriptional activating events in the former case and repressive events in the latter. Interestingly, for a smaller set of cell type-enriched genes we observed dTHSs at a given gene in both cell types, indicating regulatory activity at the gene in both cell types.

We next asked what proportion of the transcriptome differences between H and NH cells might be explained based on differential chromatin accessibility. Of the 3,282 H cell-enriched genes, 1,109 have a dTHS in one or both of the cell types, and among the 2,731 NH cell-specific genes, 724 have a dTHS in one or both cell types. Assuming that each dTHS represents a regulatory event contributing to the differential expression of its identified target gene, we could explain differential expression of 33% of the H cell-enriched genes and 27% of the NH cell-enriched genes. The remaining ~70% of the identified cell type-enriched genes without clear chromatin accessibility differences may be explained in numerous ways. These genes may not require a change in chromatin accessibility, changes in chromatin accessibility may fall below our limit of detection, or these transcripts may be primarily regulated at the post-transcriptional level rather than at the chromatin-accessibility level that we measured.

Another key question relates to the significance of the cell-type-enriched dTHSs that do not map to differentially expressed genes. These could be explained by an inability to detect all differentially expressed genes, perhaps simply due to the stringency of our definition of cell type-enriched genes. An important biological possibility to consider is that many of these regulatory regions do not in fact regulate the closest gene, but rather act over a distance such that they are orphaned from their true target genes in our analysis. Another possibility is that many of the differential protein binding events represented by these dTHSs are unrelated to transcriptional regulation.

Overall, the accessible chromatin landscapes of the root epidermal H and NH cells appear to be nearly identical in a qualitative sense, but differ significantly at several thousand sites in each cell type. The reasons for the quantitative, rather than all-or-nothing, nature of this phenomenon are not entirely clear. Are the accessibility differences between cell types reflective of unique protein assemblages at the same element in different cell types, or do they instead reflect differences in abundance of the same proteins at an element in different cell types? While these questions certainly warrant further investigation and experimentation, we can gain further insight into the regulatory differences between cell types through deeper examination of the differentially accessible chromatin regions in each.

### TF motifs in cell type-specific THSs identify regulators and their target genes

As a means of identifying specific transcription factors (TFs) that might be important in specifying the H and NH cell fates, we sought to identify overrepresented motifs in the differentially accessible regions of each cell type. We used each set of cell type-enriched dTHSs as input for MEME-ChIP analyses (Machanick and Bailey, 2011) and examined the resulting lists of overrepresented motifs. We initially found 219 motifs that were significantly overrepresented relative to genomic background only in H cell-enriched dTHSs and 12 that were significantly overrepresented only in NH cell-enriched dTHSs (Table S6). In order to narrow our list of candidate TFs to pursue, we vetted these lists of potential cell type-enriched TFs by considering their transcript levels in each cell type as well as the availability of genome-wide binding data. Based on the available data, we narrowed our search to five transcription factors of interest: four H cell-enriched TF genes (MYB33, ABI5, NAC083, and At5g04390) and one NH-enriched TF gene (WRKY27) (Table 3).

**Table 3.** Transcription factor motifs overrepresented in cell type-enriched differential transposase hypersensitive sites (dTHSs). Cell type-enriched dTHSs were analyzed for over-represented TF motifs using MEME-ChIP software, and several significantly matching factors are shown in the table. Cell specificity indicates the cell type-enriched dTHS set from which each factor was exclusively enriched, and hair/non-hair FPKM ratio indicates expression specificity of each factor using RNA-seq data from Li et al. (2016) *Developmental Cell*. Significant occurrences of each TF motif were identified across the *Arabidopsis* genome, and the percentage of these motif occurrences that fall within known binding sites for that factor (based on published ChIP-seq or DAP-seq datasets) are indicated in Column 5. Percentages are calculated by the number of motif occurrences in known binding sites/total number of motif occurrences in the genome. Column 6 indicates the percentage of THSs from the relevant cell type that contain a motif for a factor and also overlap with a known binding site for the factor (high confidence binding sites).

| Transcription Factor | Family | Cell specificity | Hair/non-hair FPKM ratio | Percentage of genomic motif occurrences in known binding sites | Percentage of motif-containing THSs that overlap a known binding site (high confidence binding sites) |
| --- | --- | --- | --- | --- | --- |
| AT5g06100 (MYB33) | MYB | Hair | 2000 | 42.2%<br>(7,038/16,655) | 69.9%<br>(1,473/2,106) |
| AT2g36270 (ABI5) | bZIP | Hair | 3.3 | 26.2%<br>(7,261/27,656) | 57.5%<br>(2,814/4,891) |
| At5g04390 | C2H2 | Hair | 17 | 9.5%<br>(3,850/40,305) | 8.5%<br>(282/3,290) |
| At5g13180 (NAC083) | NAC | Hair | 2 | 51.3%<br>(13,762/26,815) | 48.3%<br>(1,169/2,419) |
| ATgG52830 (WRKY27) | WRKY | Non-hair | 0.35 | 15.3%<br>(4,458/29,126) | 23.6%<br>(1,169/2,419) |

We next attempted to directly identify the binding sites for each TF by differential ATAC-seq footprinting between the cell types. The logic behind this approach is the same as that for DNase-seq footprinting – that the regions around a TF binding site are hypersensitive to the nuclease or transposase due to nucleosome displacement, but the sites of physical contact between the TF and DNA will be protected from transposon insertion/cutting, and thus leave behind a characteristic “footprint” of reduced accessibility on a background of high accessibility (Hesselberth et al., 2009; Vierstra and Stamatoyannopoulos, 2016). We reasoned that we could identify binding sites for each of these cell type-enriched TFs by comparing the footprint signal at each predicted binding site (a motif occurrence within a THS) between H and NH cells.

For this analysis, we examined the transposase integration patterns around the motifs of each TF in both cell types as well as in purified genomic DNA subjected to ATAC-seq, to control for transposase sequence bias. It was recently reported in *Arabidopsis* that many TF motifs exhibit conspicuous transposase integration bias on naked DNA (Lu et al., 2017), and our results were in line with these findings for all five TFs of interest here (Figure S8). While we observed footprint-like patterns in the motif-containing THSs in our ATAC-seq data, these patterns in each case were also evident on purified genomic DNA. As such, it was not possible to distinguish true binding sites from these data, as any footprint signal arising from TF binding was already obscured by the transposase integration bias. For unknown reasons, many TF motif DNA sequences seem to inherently evoke hyper- and/or hypo-integration by the transposase, and this automatically obscures any potentially informative footprint signal that could be obtained by integration during ATAC-seq on nuclei. Similar technical concerns have also been raised for DNaseI footprinting (Sung et al., 2016). These results suggest that the ATAC-seq footprinting approach may be useful for certain TFs, but these will likely need to be examined on a case-by-case basis. Given this issue and the resulting lack of evidence for footprints of our TFs of interest, we decided to take the approach of defining TF target sites as we did for our studies of root tip TFs.

As described earlier, we defined high confidence binding sites for the 5 TFs of interest as TF motif-containing THSs in the cell type of interest (predicted binding sites) that *also* overlapped with an enriched region for the TF in publicly available DAP-seq data (O'Malley et al., 2016a) or ChIP-seq data (Figure S4). Assigning these high confidence binding sites to their nearest TSS allowed us to define thousands of target genes for these factors in the root epidermal cell types (Table 3 and Table S7). Compared to our analysis of root tip TFs, our capability to predict target sites based on motif occurrences in THSs was much reduced for the four H cell-enriched and one NH cell-enriched TFs examined here. For further analyses, we decide to focus on three of the TFs that were more highly expressed in the H cell type and had the largest number of high confidence target genes: ABI5, MYB33, and NAC083.

We first asked how many of the high confidence target genes for these TFs were also preferentially expressed in one cell type or the other. We found that for all three TFs, a large percentage of the total target genes are H cell-enriched in their expression (17–21%), while many others are NH cell-enriched (6–9%) (Figure 5A). These results are intriguing as they suggest that the activities of these TFs may be generally context-dependent. At the same time however, the majority of the target genes for each TF were not more highly expressed in one cell type compared to the other.

**Figure 5.**
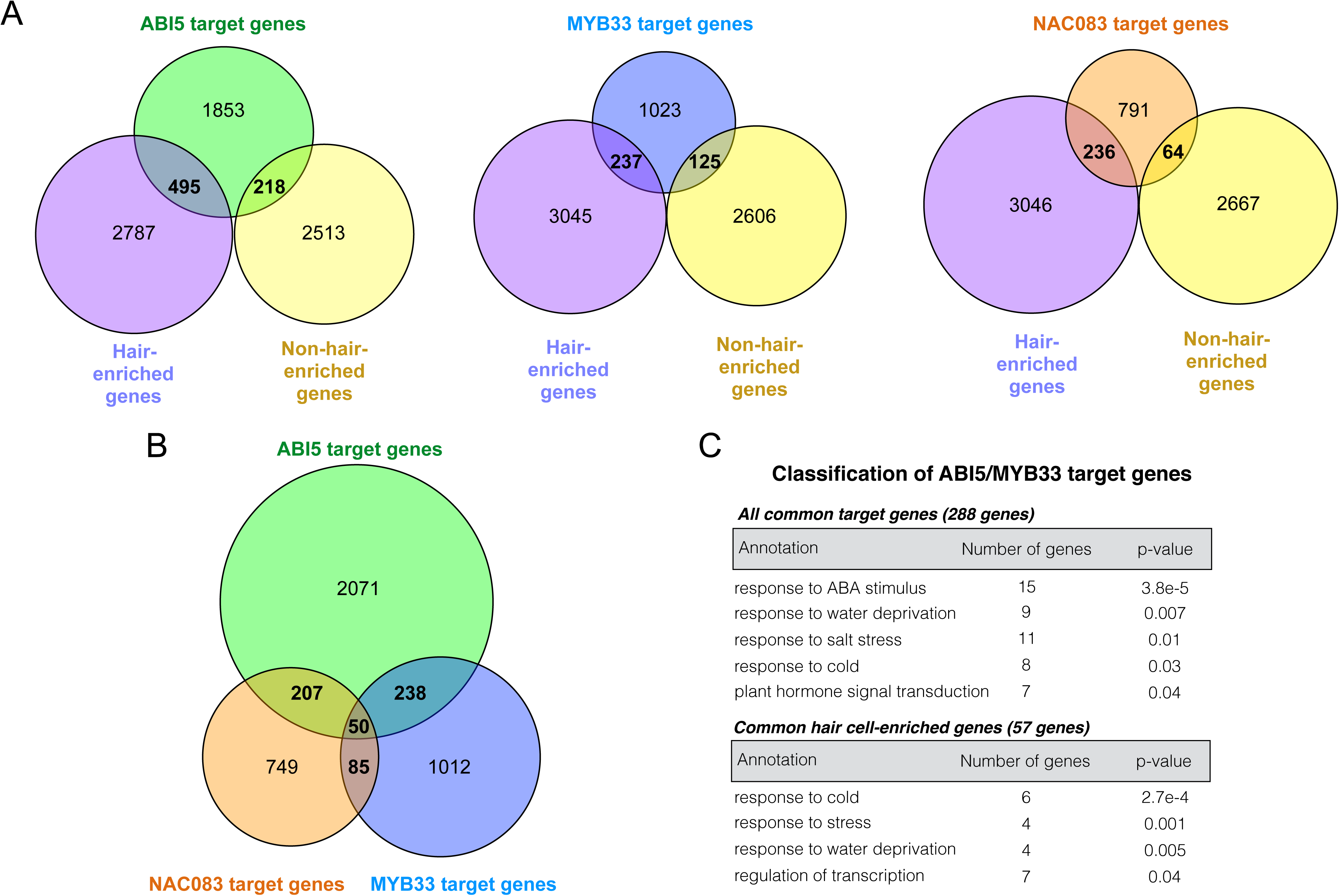
Targeting of cell type-enriched genes by H cell-enriched TFs, and co-regulatory associations among H cell-enriched TFs. Genome-wide high confidence binding sites for each TF were defined as open chromatin regions in the hair cell that contain a significant motif occurrence for the factor and also overlap with a known enriched region for that factor from DAP-seq or ChIP-seq data. Target genes were defined by assigning each high confidence binding site to the nearest TSS. **(A)** Venn diagrams showing high confidence target genes for ABI5, MYB33, and NAC083 and their overlap with cell type-enriched genes. **(B)** Overlap of ABI5, MYB33, and NAC083 high confidence target genes. **(C)** Gene Ontology (GO) analysis was performed to illuminate biological functions of genes co-targeted by ABI5 and MYB33. The upper panel shows significantly enriched GO terms for all 288 genes targeted by both ABI5 and MYB33. For each enriched annotation term, the number of genes in the set with that term is shown, followed by the FDR-corrected p-value. The lower panel lists significantly enriched GO-terms for the 57 hair cell-enriched genes co-targeted by ABI5 and MYB33. The seven hair cell-enriched genes associated with the term *regulation of transcription* were chosen for further analysis. All annotation terms in the lists are at the Biological Process level except for the KEGG pathway term ‘plant hormone signal transduction’.

Each of these H cell-enriched TFs could activate other H cell-enriched genes, but what are their functions at regulatory elements near genes that are expressed at low levels in the H cell and high levels in the NH cell? One possibility is that these factors are activators of transcription in the context of H cell-enriched genes but act as repressors or are neutral toward the target genes that are NH cell-enriched in their expression. This may reflect context-dependency in the sense that the effect on transcription of a target gene may depend on the local milieu of other factors.

We next examined whether ABI5, MYB33, and NAC083 target any of the same genes. Similar to the root tip TFs examined previously, we found that these three TFs also appear to have extensive co-regulatory relationships (Figure 5B). For example, 207 target genes were shared between ABI5 and NAC083, 238 were shared between ABI5 and MYB33, and 50 target genes were shared by all 3 factors. We further analyzed the genes that were co-targeted by ABI5 and MYB33, finding that 57 of the co-targeted genes were H-cell enriched. As such, we performed Gene Ontology (GO) analysis on the H cell-enriched targets as well as the full set of target genes to gain insight into the functions of this co-regulatory relationship (Figure 5C). Many of the ABI5/MYB33 target genes were annotated as being involved in responses to ABA as well as water, salt, and cold stress. This is consistent with the known roles of these proteins in ABA signaling (Finkelstein and Lynch, 2000; Reyes and Chua, 2007). Interesting, seven of the 57 ABI5/MYB33 target genes that were H cell-enriched were also annotated with the term *regulation of transcription*, suggesting that ABI5 and MYB33 may be at the apex of a transcriptional regulatory cascade in the H cell type.

### Identification of a new regulatory module in the root hair cell type

Based on our findings that ABI5 and MYB33 co-target seven H cell-enriched TFs, we decided to investigate this potential pathway further. Among the seven TFs putatively co-regulated by ABI5 and MYB33 and having H cell-enriched transcript expression were DEAR5, ERF11, At3g49930, SCL8, NAC087, and two additional MYB factors: MYB44 and MYB77. Aside from MYB77, none of these TFs had been previously reported to produce root-specific phenotypes when mutated. MYB77 was previously shown to interact with Auxin Response Factors (ARFs) (Shin et al., 2007) and to be involved in lateral root development through promotion of auxin-responsive gene expression (Shin et al., 2007). Interestingly, the ABA receptor, PYL8, was shown to physically interact with both MYB77 and MYB44, and to promote auxin-responsive transcription by MYB77 (Zhao et al., 2014). MYB44 has also been implicated in ABA signaling through direct interaction with an additional ABA receptor, PYL9 (Li et al., 2014), as well as repression of jasmonic acid (JA)-responsive transcription (Jung et al., 2010). These factors have additionally been implicated in salicylic acid (SA) and ethylene signaling (Yanhui et al., 2006; Shim et al., 2013). Given that MYB44 and MYB77 are paralogs (Dubos et al., 2010) that appear to integrate multiple hormone response pathways in a partly redundant manner (Jaradat et al., 2013), we decided to identify high confidence target genes (Figure S4) for each of them for further study.

We again defined high confidence binding sites as THSs in H cells that contain a significant motif occurrence for the factor and also overlap with a DAP-seq or ChIP-seq enriched region for that factor. Using this approach, we found that MYB44 and MYB77 each target over 1,000 genes individually and co-target 483 genes (Figure 6A). In addition, MYB44 and MYB77 appear to regulate one another, while MYB77 also appears to target itself. This feature of self-reinforcing co-regulation could serve as an amplifying and sustaining mechanism to maintain the activity of this module once activated by ABI5, MYB33, and potentially other upstream factors.

**Figure 6.**
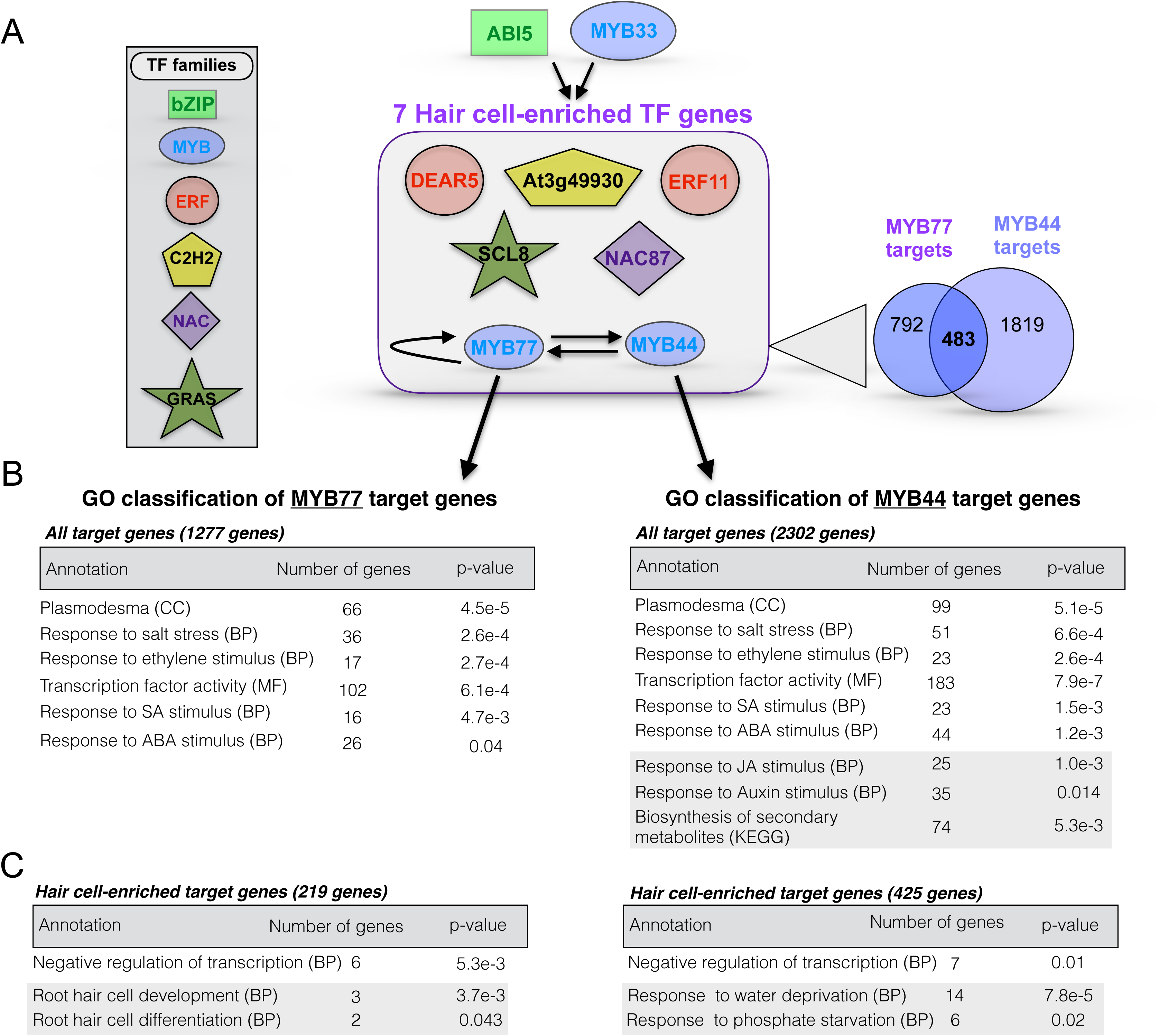
A transcriptional regulatory module in the root hair cell type. **(A)** Diagram of the proposed regulatory module under control of ABI5 and MYB33. As referenced in **Figure 5C**, ABI5 and MYB33 co-target seven TFs that are preferentially expressed in the hair cell relative to the non-hair cell type. The family classification of each of the seven TFs is denoted in the figure key. Among the seven hair cell-specific target TFs are two MYB family members, MYB77 and MYB44. High confidence binding sites for these two MYB factors were again defined as open chromatin regions in the hair cell that contain a significant motif occurrence for the factor and also overlap with a known enriched region for that factor from DAP-seq or ChIP-seq data. Each high confidence binding site was then assigned to the nearest TSS to define the target gene for that site. This analysis revealed that MYB44 and MYB77 target each other, and MYB77 targets itself. Both factors target thousands of additional genes, 483 of which are in common (Venn diagram on the lower right of the schematic). Arrows coming down from MYB77 and MYB44 point to GO analyses of that factor’s target genes. **(B)** The upper tables represent enriched annotation terms for all target genes of the factor, regardless of differential expression between H and NH cells, while the lower tables **(C)** represent enrichment of terms within target genes that are preferentially expressed in the hair cell relative to the non-hair cell. Annotation term levels are indicated as Cellular Component (CC), Biological Process (BP), Molecular Function (MF) or KEGG pathway (KEGG). For each annotation, the number of target genes associated with that term is shown to the right of the term, followed by the FDR-corrected p-value for the term enrichment in the rightmost column. Groups of terms boxed in gray are those that differ between MYB44 and MYB77. The structure of the module suggests that ABI5 and MYB33 drive a cascade of TFs including MYB77 and MYB44, which act to amplify this signal and also further regulate many additional TFs. Additional target genes of MYB77 and MYB44 include hair cell differentiation factors, hormone response genes, secondary metabolic genes, and genes encoding components of important cellular structures such as plasmodesmata.

To gain a deeper understanding of the impact of MYB44 and MYB77 on downstream processes, we performed Gene Ontology (GO) analysis of the target genes for each factor. First considering all target genes, regardless of their expression in the H cell type, we found a variety of overrepresented GO terms for each that were consistent with the known roles of these factors in hormone signaling (Figure 6B). For example, both factors targeted a large number of genes annotated with the terms *response to ABA stimulus*, *response to ethylene stimulus*, and *response to SA stimulus*. Additionally, MYB44 alone targeted many genes with the annotation *response to JA stimulus*, consistent with its previously reported role as a negative regulator of JA signaling (Jung et al., 2010). Interestingly, the largest overrepresented gene functional category for both factors was *transcription factor activity* (102 genes for MYB77 and 183 genes for MYB44). This indeed further suggests that these factors initiate a cascade of transcriptional effects. The next-largest overrepresented term was *plasmodesma*, indicating that production and/or regulation of cell-cell connecting structures are likely controlled by these factors. Plasmodesmata are important for numerous epidermal functions including cell-to-cell movement of TFs such as CPC and TRY (Schellmann et al., 2002; Wada et al., 2002) and transport of other macromolecules and metabolites (Lucas and Lee, 2004).

We also analyzed overrepresented ontology terms in the MYB77 and MYB44 targets that were classified as H cell-enriched genes. Among the MYB77 target genes in this category were known regulators of H cell fate, while numerous H cell-enriched MYB44 target genes were annotated as being involved in response to water and phosphate starvation (Figure 6C). The ontology category that was overrepresented in both target lists was *negative regulation of transcription* (6 MYB77 targets and 7 MYB44 targets), suggesting that these factors exert additional specific effects on the H cell transcriptome by regulating a subset of potentially repressive TFs.

The fact that MYB77 and MYB44 target a large number of genes that show H cell-enriched expression suggests that these factors serve as activators of transcription at these targets, and this is supported by published accounts of transcriptional control by these factors (Persak and Pitzschke, 2014). However, both factors also target NH cell-enriched genes as well as genes without preferential expression between the cell types. This phenomenon was also observed for the H-enriched TFs ABI5, MYB33, and NAC083 (Figure 5), suggesting that certain TFs may generally serve as activators but may also have context-dependent repressive functions. Such a functional switch could occur through direct mechanisms such as structural alteration by alternative splicing or post-translational modification, functional alteration by partnering with a specific TF or chromatin-modifying complex, or perhaps indirectly by binding to a target site to occlude the binding of other factors necessary for transcriptional activation. The numerous reports of dual function transcription factors in animals and plants support the notion that this may be a general phenomenon (Ikeda et al., 2009; Boyle and Despres, 2010; Li et al., 2016b).

Collectively these results suggest that the MYB44/MYB77 module in the H cell specifies a cascade of downstream transcriptional regulation, some of which is positive and some of which is negative. This module likely represents an important hub in controlling H cell fate as well as a variety of physiological functions and environmental responses in this cell type. The fact that MYB77 was also discovered in our analyses of root tip TFs suggests that this factor likely has a broader role in other cell types during early root development, in addition to a role in specification of the H cell versus the NH cell fate. An important next step will be to perform genetic manipulations of these factors (knockout and inducible overexpression, for example), in order to test and elaborate on the specific predictions made by our model.

## Summary and conclusions

In this study, we used ATAC-seq profiling of accessible chromatin to investigate questions regarding the transcriptional regulatory landscape of plant genomes and its conservation across species. We also investigated the similarities and differences in open chromatin landscapes in two root cell types that arise from a common progenitor, allowing us to identify and analyze TFs that act specifically in one cell type versus the other. Overall, we are able to gain several new insights from this work.

In optimization of our ATAC-seq procedures, we found that the assay can be performed effectively on crudely purified nuclei but that this approach is limited by the large proportion of reads arising from organelle genomes (Table 1). This issue is ameliorated by the use of the INTACT system to affinity-purify nuclei for ATAC-seq, which also provides access to individual cell types. Consistent with previous reports, we found that the data derived from ATAC-seq are highly similar to those from DNase-seq (Figure 1). In comparing our root tip ATAC-seq data to DNase-seq data from whole roots, we found that some hypersensitive regions were detected in one assay but not the other. This discrepancy is most likely attributable to differences in starting tissue and laboratory conditions, rather than biological differences in the chromatin regions sensitive to DNaseI versus the hyperactive Tn5 transposase. This interpretation would fit with the large number of differences also observed in THS overlap between *Arabidopsis* root tip and epidermal cell types.

In a comparison of open chromatin among the root tip epigenomes of *Arabidopsis*, *Medicago*, tomato, and rice, we found the genomic distribution of THSs in each were highly similar. About 75% of THSs lie outside of transcribed regions, and the majority of these THSs are found within 3 kb upstream of the TSS in all species (Figure 2). Thus, the distance of upstream THSs from the TSS is relatively consistent among species and is not directly proportional to genome size or intergenic space for these representative plant species. Among genes with an upstream THS, 70% of these genes in *Arabidopsis*, *Medicago*, and rice have a single such feature, 20% have two upstream THSs, and less than 10% have three or more. In contrast, only 27% of tomato genes with an upstream THS have a single THS, 20% have two, and the proportion with 4–10 THSs is 2–7 times higher than that for any other species examined. This increase in THS number in tomato could be reflective of an increase in the number of regulatory elements per gene, but is perhaps more likely a result of the greater number of long-terminal repeat retrotransposons near genes in this species (Xu and Du, 2014). In either case, our investigation revealed that open chromatin sites – and by extension transcriptional regulatory elements – in all four species are focused in the TSS-proximal upstream regions and are relatively few in number per gene. This suggests that transcriptional regulatory elements in plants are generally fewer in number and are closer to the genes they regulate than those of animal genomes. For example, the median distance from an enhancer to its target TSSs in *Drosophila* was found to be 10 kb, and it was estimated that each gene had an average of four enhancers (Kvon et al., 2014). It was also recently reported that in human T cells, the median distance between enhancers and promoters was 130 kb, far greater than the distances we have observed here across plant species (Mumbach et al., 2017).

Analysis of over-represented TF motifs in THSs across species suggested that many of the same TFs are at play in early root development in all species. Perhaps more surprisingly, co-regulation of specific gene sets by multiple TFs seems to be frequently maintained across species (Figure 3). Taken together with the lack of shared open chromatin profiles among orthologous genes and expressologs, these findings suggest that transcriptional regulatory elements may relocate over evolutionary time within a window of several kilobases upstream of the TSS, but regulatory control by specific TFs is relatively stable.

Our comparison of the two *Arabidopsis* root epidermal cell types, the hair (H) and non-hair (NH) cells, revealed that open chromatin profiles were highly similar between cell types. By examining THSs that were exclusive to one cell type, we were able to find several thousand THSs that were quantitatively more accessible in each cell type compared to the other (Figure 4). Mapping of these differential THSs (dTHSs) to their nearest genes revealed that in each cell type there were many dTHSs that were near genes expressed more abundantly in that cell type, as well as many near genes with the opposite expression pattern. This suggests that some dTHSs represented transcriptional activating events whereas others were repressive in nature.

Analysis of TF motifs at these dTHSs between cell types identified a suite of TFs that were more highly expressed in H cells and whose motifs were significantly overrepresented in H cell-enriched dTHSs. Analysis of three of these TFs – ABI5, MYB33, and NAC083 – revealed that each factor targets a large number of H cell-enriched genes as well as a smaller number of NH cell-enriched genes (Figure 5). These factors also have many overlapping target genes among them, and ABI5 and MYB33 both target seven additional H cell-enriched TFs. Among these seven H-enriched TFs are two additional MYB factors: MYB77 and MYB44 (Figure 6). Examination of the high confidence target genes of MYB77 and MYB44 revealed that these paralogous factors appeared to regulate each other as well as many other common target genes, including large numbers of other TF genes. Hundreds of the MYB77 and MYB44 target genes were also more highly expressed in the H cell relative to the NH cell, suggesting that these factors set off a broad transcriptional cascade in the H cell type. In addition, they appear to directly regulate many H cell-enriched genes involved in cell fate specification and water and phosphate acquisition. This type of cooperative action by pairs of MYB paralogs has also been documented recently in *Arabidopsis* and other species (Millar and Gubler, 2005; Matus et al., 2017; Wang et al., 2017), and the fact that many target genes for each MYB factor are not regulated by the other may reflect a degree of subfunctionalization between the paralogs.

An important question arising from our results is whether classifying a TF as strictly an activator or repressor is generally accurate in most cases. For example, the H cell-enriched TFs that we examined all have apparent target genes that are highly expressed in the H cell type as well as targets that are expressed at very low levels, if at all, in the H cell type. In fact, these latter genes are often much more highly expressed in the NH cell type. Given that a number of these TFs have been shown to activate transcription in specific cases, this suggests that they promote the transcription of H cell-enriched targets and either repress or have no effect on NH cell-enriched target genes. One explanation for this phenomenon is that these TFs have “dual functionality” as activators and repressors, depending on the context (Bauer et al., 2010). However, it is equally possible that these factors do not play a direct role in gene repression. For example, the binding of an activator near a repressed gene may be functionally irrelevant to the regulation of that gene, or it may be the case that other gene-specific repressors may also be bound nearby and override the activity of the activator. This phenomenon will be worth exploring as it may deepen our understanding of the intricacies of transcriptional control.

In this study, we outline a widely applicable approach for combining chromatin accessibility profiling with available genome-wide binding data to construct models of TF regulatory networks. The putative TF regulatory pathways we have illuminated through our comparison across species and cell types provide important hypotheses regarding the evolution of gene regulatory mechanisms in plants and the mechanisms of cell fate specification, that are now open to experimental analysis.

## METHODS

### Plant materials and growth conditions

Plants used in this study were of the *Arabidopsis thaliana* Col-0 ecotype, the A17 ecotype of *Medicago truncatula*, the M82 LA3475 cultivar of tomato (*Solanum lycopersicum*), and the Nipponbare cultivar of rice (*Oryza sativa*). Transgenic plants of each species for INTACT were produced by transformation with a binary vector carrying both a constitutively expressed biotin ligase and constitutively expressed nuclear tagging fusion protein (NTF) containing a nuclear outer membrane association domain (Ron et al., 2014). The binary vector used for *Medicago* was identical to the tomato vector (Ron et al., 2014), but was constructed in a pB7WG vector containing phosphinothricin resistance gene for plant selection and it retains the original *AtACT2p* promoter. The binary vector used for rice is described in Reynoso *et al.* (submitted). Transformation of rice was carried out at UC Riverside and tomato transformation was carried out at the UC Davis plant transformation facility. *Arabidopsis* plants were transformed by the floral dip method (Clough and Bent, 1998) and composite transgenic *Medicago* plants were produced according to established procedures (Limpens et al., 2004).

For root tip chromatin studies, constitutive INTACT transgenic plant seeds were surface sterilized and sown on ½-strength Murashige and Skoog (MS) media (Murashige and Skoog, 1962) with 1% (w/v) sucrose in 150 mm diameter Petri plates, except for tomato and rice where full-strength MS with 1% (w/v) sucrose and without vitamins was used. Seedlings were grown on vertically oriented plates in controlled growth chambers for 7 days after germination, at which point the 1 cm root tips were harvested and frozen immediately in liquid N_2_ for subsequent nuclei isolation. The growth temperature and light intensity was 20°C and 200 µmol/m^2^/sec for *Arabidopsis* and *Medicago*, 23°C and 80 µmol/m^2^/sec for tomato, and 28°C/25°C day/night and 110 µmol/m^2^/sec for rice. Light cycles were 16 h light/8 h dark for all species.

For studies of the *Arabidopsis* root hair and non-hair cell types, previously described INTACT transgenic lines were used (Deal and Henikoff, 2010). These lines are in the Col-0 background and carry a constitutively expressed biotin ligase gene (*ACT2p:BirA*) and a transgene conferring cell type-specific expression of the NTF gene (from the *GLABRA2* promoter in non-hair cells or the *ACTIN DEPOLYMERIZING FACTOR8* promoter in root hair cells). Plants were grown vertically on plates as described above for 7 days, at which point 1.25 cm segments from within the fully differentiated cell zone were harvested and flash frozen in liquid N_2_. This segment of the root contains only fully differentiated cells and excludes the root tip below and any lateral roots above.

### Nuclei isolation

For comparison of ATAC-seq using crude and INTACT-purified *Arabidopsis* nuclei, a constitutive INTACT line was used (*ACT2p:BirA*/*UBQ10p:NTF*) (Sullivan et al., 2014) and nuclei were isolated as described previously (Bajic et al., 2017). In short, after growth and harvesting as described above, 1–3 g of root tips were ground to a powder in liquid N_2_ in a mortar and pestle and then resuspended in 10 ml of NPB (20 mM MOPS [pH 7], 40 mM NaCl, 90 mM KCl, 2 mM EDTA, 0.5 mM EGTA, 0.5 mM spermidine, 0.2 mM spermine, 1× Roche Complete protease inhibitors) with further grinding. This suspension was then filtered through a 70 µM cell strainer and centrifuged at 1,200 x g for 10 min at 4° C. After decanting, the nuclei pellet was resuspended in 1 ml of NPB and split into two 0.5 ml fractions in new tubes. Nuclei from one fraction were purified by INTACT using streptavidin-coated magnetic beads as previously described (Bajic et al., 2017) and kept on ice prior to counting and subsequent transposase integration reaction. Nuclei from the other fraction were purified by non-ionic detergent lysis of organelles and sucrose sedimentation, as previously described (Bajic et al., 2017). Briefly, these nuclei in 0.5 ml of NPB were pelleted at 1,200 x g for 10 min at 4° C, decanted, and resuspended thoroughly in 1 ml of cold EB2 (0.25 M sucrose, 10 mM Tris [pH 8], 10 mM MgCl_2_, 1% Triton X-100, and 1× Roche Complete protease inhibitors). Nuclei were then pelleted at 1,200 x g for 10 min at 4° C, decanted, and resuspended in 300 µl of EB3 (1.7 M sucrose, 10 mM Tris [pH 8], 2 mM MgCl_2_, 0.15% Triton X-100, and 1× Roche Complete protease inhibitors). This suspension was then layered gently on top of 300 µl of fresh EB3 in a 1.5 ml tube and centrifuged at 16,000 x g for 10 minutes at 4° C. Pelleted nuclei were then resuspended in 1 ml of cold NPB and kept on ice prior to counting and transposase integration.

For INTACT purification of total nuclei from root tips of *Medicago*, tomato and rice, as well as purification of *Arabidopsis* root hair and non-hair cell nuclei, 1–3 g of starting tissue was used. In all cases, nuclei were purified by INTACT and nuclei yields were quantified as described previously (Bajic et al., 2017).

### Assay for transposase-accessible chromatin with sequencing (ATAC-seq)

Freshly purified nuclei to be used for ATAC-seq were kept on ice prior to the transposase integration reaction and never frozen. Transposase integration reactions and sequencing library preparations were then carried out as previously described (Bajic et al., 2017). In brief, 50,000 purified nuclei or 50 ng of *Arabidopsis* leaf genomic DNA were used in each 50 µl transposase integration reaction for 30 min at 37° C using Nextera reagents (Illumina, FC-121-1030). DNA fragments were purified using the Minelute PCR purification kit (Qiagen), eluted in 11 µl of elution buffer, and the entirety of each sample was then amplified using High Fidelity PCR Mix (NEB) and custom barcoded primers for 9–12 total PCR cycles. These amplified ATAC-seq libraries were purified using AMPure XP beads (Beckman Coulter), quantified by qPCR with the NEBNext Library Quantification Kit (NEB), and analyzed on a Bioanalyzer High Sensitivity DNA Chip (Agilent) prior to pooling and sequencing.

### High throughput sequencing

Sequencing was carried out using the Illumina NextSeq 500 or HiSeq2000 instrument at the Georgia Genomics Facility at the University of Georgia. Sequencing reads were either single-end 50 nt or paired-end 36 nt and all libraries that were to be directly compared were pooled and sequenced on the same flow cell.

### Sequence read mapping, processing, and visualization

Sequencing reads were mapped to their corresponding genome of origin using Bowtie2 software (Langmead and Salzberg, 2012) with default parameters. Genome builds used in this study were *Arabidopsis* version TAIR10, *Medicago* version Mt4.0, Tomato version SL2.4, and Rice version IRGSP 1.0.30. Mapped reads in *.sam* format were converted to *.bam* format and sorted using Samtools 0.1.19 (Li et al., 2009). Mapped reads were then filtered using Samtools to retain only those reads with a mapping quality score of 2 or higher (Samtools “*view”* command with option “*-q 2”* to set mapping quality cutoff). *Arabidopsis* ATAC-seq reads were further filtered with Samtools to remove those mapping to either the chloroplast or mitochondrial genomes, and root hair and non-hair cell datasets were also subsampled such that the experiments within a biological replicate had the same number of mapped reads prior to further analysis. For normalization and visualization, the filtered, sorted *.bam* files were converted to bigwig format using the “*bamcoverage”* script in deepTools 2.0 (Ramirez et al., 2016) with a bin size of 1 bp and RPKM normalization. Use of the term *normalization* in this paper refers to this process. Heatmaps and average plots displaying ATAC-seq data were also generated using the “*computeMatrix”* and “*plotHeatmap”* functions in the deepTools package. Genome browser images were made using the Integrative Genomics Viewer (IGV) 2.3.68 (Thorvaldsdottir et al., 2013) with bigwig files processed as described above.

### Identification of orthologous genes among species

Orthologous genes among species were selected exclusively from syntenic regions of the four genomes. Syntenic orthologs were identified using a combination of CoGe SynFind (https://genomevolution.org/CoGe/SynFind.pl) with default parameters, and CoGe SynMap (https://genomevolution.org/coge/SynMap.pl) with QuotaAlign feature selected and a minimum of six aligned pairs required (Lyons and Freeling, 2008; Lyons et al., 2008).

### Peak calling to detect transposase hypersensitive sites (THSs)

Peak calling on ATAC-seq data was performed using the “*Findpeaks”* function of the HOMER package (Heinz et al., 2010). The parameters “*-region”* and “-*minDist 150”* were used to allow identification of variable length peaks and to set a minimum distance of 150 bp between peaks before they are merged into a single peak, respectively. We refer to the peaks called in this way as “transposase hypersensitive sites”, or THSs.

### Genomic distribution of THSs

For each genome, the distribution of THSs relative to genomic features was assessed using the PAVIS web tool (Huang et al., 2013) with “upstream” regions set as the 2,000 bp upstream of the annotated transcription start site and “downstream” regions set as 1,000 bp downstream of the transcript termination site.

### Transcription factor motif analyses

ATAC-seq transposase hypersensitive sites (THSs) that were found in two replicates of each sample were used for motif analysis. The regions were adjusted to the same size (500 bp for root tip THSs or 300 bp for cell type-specific dTHSs). The MEME-ChIP pipeline (Machanick and Bailey, 2011) was run on the repeat-masked fasta files representing each THS set to identify overrepresented motifs, using default parameters. For further analysis, we used the motifs derived from the DREME, MEME, and CentriMo programs that were significant matches (E value < 0.05) to known motifs. Known motifs from both Cis-BP (Weirauch et al., 2014b) and the DAP-seq database (O’Malley et al., 2016) were used in all motif searches.

### Assignment of THSs to genes

For each ATAC-seq data set the THSs were assigned to genes using the “*TSS”* function of the PeakAnnotator 1.4 program (Salmon-Divon et al., 2010). This program assigns each peak/THS to the closest transcription start site (TSS), whether upstream or downstream, and reports the distance from the peak center to the TSS based on the genome annotations described above.

### ATAC-seq footprinting

To examine motif-centered footprints for TFs of interest we used the “*dnase_average_profile.py”* script in the pyDNase package (Piper et al., 2013). The script was used in ATAC-seq mode [“*-A”* parameter] with otherwise default parameters.

### Publicly available DNase-seq, DAP-seq, ChIP-seq, and RNA-seq data

For comparison to our ATAC-seq data from root tips, we used a published DNase-seq dataset from 7-day-old whole *Arabidopsis* roots (SRX391990), which was generated from the same INTACT transgenic line used in our experiments (Sullivan et al., 2014).

Publicly available ChIP-seq and DAP-seq datasets were also used to identify genomic binding sites for transcription factors of interest. These include ABF3 (AT4G34000; SRX1720080) and MYB44 (AT5G67300; SRX1720040)(Song et al., 2016), HY5 (AT5G11260; SRX1412757), CBF2 (AT4G25470; SRX1412036), MYB77 (AT3G50060; SRX1412453), ABI5 (AT2G36270; SRX670505), MYB33 (AT5G06100; SRX1412418), NAC083 (AT5G13180; SRX1412546), MYB77 (AT3G50060; SRX1412453), WRKY27 (AT5G52830; SRX1412681), and At5g04390 (SRX1412214) (O’Malley et al., 2016). Raw reads from these files were mapped and processed as described above for ATAC-seq data, including peak calling with the HOMER package.

Published RNA-seq data from *Arabidopsis* root hair and non-hair cells (Li et al., 2016a) were used to define transcripts that were specifically enriched in the root hair cell relative to the non-hair cell (hair cell enriched genes), and vice versa (non-hair enriched genes). We defined cell type-enriched genes as those whose transcripts were at least two-fold more abundant in one cell type than the other and had an abundance of at least five RPKM in the cell type with higher expression.

### Defining high confidence target sites for transcription factors

We used FIMO (Grant et al., 2011) to identify motif occurrences for TFs of interest, and significant motif occurrences were considered to be those with a p-value < 0.0001. Genome-wide high confidence binding sites for a given transcription factor were defined as transposase hypersensitive sites in a given cell type or tissue that also contain a significant motif occurrence for the factor and also overlap with a known enriched region for that factor from DAP-seq or ChIP-seq data (see also Figure S2 for a schematic diagram of this process).

### Gene ontology analysis

Gene Ontology (GO) analyses using only *Arabidopsis* genes were carried out using the GeneCodis 3.0 program (Nogales-Cadenas et al., 2009; Tabas-Madrid et al., 2012). Hypergeometric tests were used with p-value correction using the false discovery rate (FDR) method. AgriGO was used for comparative GO analysis of gene lists among species, using default parameters (Du et al., 2010; Tian et al., 2017).

### Accession Numbers

The raw and processed ATAC-seq data described here have been deposited to the NCBI Gene Expression Omnibus (GEO) database under record number GSE101482. The characteristics of each dataset (individual accession number, read numbers and mapping characteristics, and THS statistics) are included in Table S8.

## ACKNOWLEDGEMENTS

We thank Paja Sijacic and Shannon Torres for constructive criticism of the manuscript. This work was supported by funding from the National Science Foundation (Plant Genome Research Program grant #IOS-123843) to J.B-S., N.S., S.M.B., and R.B.D.; D.A.W. was supported in part by funding from the Elise Taylor Stocking Memorial Fellowship, and K.K. was supported in part by the Finnish Cultural Foundation.

## AUTHOR CONTRIBUTIONS

R.B.D., S.M.B., N.S., J.B-S., K.A.M., M. B., K.K, and M.R, G.P., and D.A.W. designed the research project. K.A.M. performed all experiments on *Arabidopsis* root tips as well as hair and non-hair cells. M.B. performed all experiments on *Medicago* root tips, K.K., D.A.W, and K.Z. performed all experiments on tomato root tips, and M.R. and G.P. performed all experiments on rice root tips. M.W. performed all analyses of syntenic regions and identification of orthologous genes among species. K.B., M.D., and C.Q. analyzed ATAC-seq data sets with Hotspot software and also contributed expertise in other analyses. R.B.D, K.A.M, and M.B. analyzed the data, and R.B.D. drafted the manuscript with subsequent input and editing from all authors.

